# Estimating individual level plant traits at scale

**DOI:** 10.1101/556472

**Authors:** Sergio Marconi, Sarah J. Graves, Ben. G. Weinstein, Stephanie Bohlman, Ethan P. White

## Abstract

Functional ecology has increasingly focused on describing ecological communities based on their traits (measurable features affecting individuals fitness and performance). Analyzing trait distributions within and among forests could significantly improve understanding of community composition and ecosystem function. Historically, data on trait distributions are generated by (1) collecting a small number of leaves from a small number of trees, which suffers from limited sampling but produces information at the fundamental ecological unit (the individual); or (2) using remote sensing images to infer traits, producing information continuously across large regions, but as plots (containing multiple trees of different species) or pixels, not individuals. Remote sensing methods that identify individual trees and estimate their traits would provide the benefits of both approaches, producing continuous large-scale data linked to biological individuals. We used data from the National Ecological Observatory Network (NEON) to develop a method to scale up functional traits from 160 trees to the millions of trees within the spatial extent of two NEON sites. The pipeline consists of three stages: 1) image segmentation, to identify individual trees and estimate structural traits; 2) ensemble of models to infer leaf mass area (LMA), nitrogen, carbon, and phosphorus content using hyperspectral signatures, and DBH from allometry; and 3) predictions for segmented crowns for the full remote sensing footprint at the NEON sites.

The R^2^ values on held out test data ranged from 0.41 to 0.75 on held out test data. The ensemble approach performed better than single partial least squares models. Carbon performed poorly compared to other traits (R^2^ of 0.41). The crown segmentation step contributed the most uncertainty in the pipeline, due to over-segmentation. The pipeline produced good estimates of DBH (R^2^ of 0.62 on held out data). Trait predictions for crowns performed significantly better than comparable predictions on pixels, resulting in improvement of R^2^ on test data of between to 0.26. We used the pipeline to produce individual level trait data for ∼5 million individual crowns, covering a total extent of ∼360 km^2^. This large dataset allows testing ecological questions on landscape scales, revealing that foliar traits are correlated with structural traits and environmental conditions.

## 1. Introduction

Functional traits are biochemical, physiological and structural characters that influence organism performance or fitness (Nock et al., 2016). They are central to how organisms perform under different environmental conditions, interact with other species, and influence the ecosystems to which they belong (McGill 2006, Dwyer et al., 2017, Collalti et al., 2019). For individual organisms, traits influence core demographic parameters including survival and reproduction. At the species level, traits influence species distributions and how species respond to changes in land use and climate (Pollock et al., 2012). At the ecosystem level, organismal traits influence biogeochemical cycles and habitat availability for other species (e.g., Fisichelli et al., 2015). Given their central importance across multiple levels of organization, understanding how traits vary within and among species, across environmental gradients, and through time is essential to understanding many areas of ecology and predicting how ecological systems will change in the future (McGill 2006, Lawler et al. 2010, Valladares et al., 2014, Diaz et al., 2016).

In trees, two commonly studied groups of traits are specific to (1) properties of leaves (e.g., leaf mass per area, nitrogen and phosphorus concentration) and (2) the size structure of the full tree (e.g. height, dbh, canopy size). These characters hold different information about tree properties and how they link to forest functions. Nitrogen and phosphorus, for example, are fundamental proxies of leaf productivity because of their fundamental role in photosynthesis (Tang et al., 2018); LMA is a widely used indicator of different leaf anatomy and foliar structure strategies (Poorter et al., 2008); and tree height and dbh are indicators of tree structure and growth. Having access to measures of both leaf and structural (or physiognomic) traits for individual trees across the landscape potentially unlocks the ability to explore different dimensions of biodiversity together, investigate how these properties influence each other and affect competition among neighboring trees, and link to tree functions like growth and carbon exchange. However, exploring the links between leaf and structural traits across landscape is challenging, in part because of the differences in the design of their sampling approach.

Historically, structural traits are collected for thousands of trees in targeted areas via programs such as the US Forest Inventory and Analysis, whereas studies of leaf chemical traits have relied on collecting samples of a few leaves from a small number of individuals. These values are used to estimate the average trait values for each species and to explore how ecosystem level leaf traits vary biogeographically or through time by assuming that all individuals of a species in a region share the same trait value (Swenson et al., 2010, Clark et al., 2016). This approach is necessary because it is expensive and time consuming to collect individual level leaf trait data, but it fails to describe trait variation within species driven by evolution and plastic responses to the conditions an individual or population experiences (Messier et al., 2017, Niinemets et al, 2017, Muller et al. 2010, Nicotra et al. 2010, Albert et al. 2010, Callaway et al. 2003). Moreover, since the number of leaf trait records is often orders of magnitude smaller than tree structural trait records, discrepancy in their sample size may affect the generality of relationships observed between leaf and structural traits at landscape scales. This limitation is magnified when studying changing environments (across space or time) because of bias in where the data for each species is collected. Data are typically collected in small subsets of the full range of conditions that species experience and are often selected in a biased manner that fits the purpose of the original studies (e.g., selecting individuals of a particular health status, size or species). Measuring traits systematically across geographic gradients would address this limitation, but is not feasible with traditional field methods (Anderson-Teixeira 2015).

An alternative approach that allows continuous estimation of traits across the landscape is to use remote sensing data (Kerr & Ostrovsky, 2003, Homolova et al. 2013, Houborg et al., 2015). For example, (a) hyperspectral remote sensing imagery is used to estimate the chemical composition of sunlit leaves by measuring light absorption and reflectance in the visible and near-infrared spectrum (Asner et al., 2017), and (b) light detection and ranging (LiDAR) is used to measure vertical and horizontal vegetation structure (Andersen et al., 2005). Leveraging remote sensing approaches allows for measuring trait information continuously at landscape scales. Together, LiDAR and hyperspectral data can be used to estimate many of the standard leaf and structural tree traits for trees (Serbin et al., 2014, Singh et al., 2015, Asner et al., 2017, Barbosa et al., 2017).

Traditionally, remote sensing applications use either the pixel (the smallest resolution component of the image, Audebert et al., 2019) or the plot (a region of space typically containing multiple individuals, Singh et al., 2015,) as the fundamental unit. This is a natural result of the structure of the remote sensing data combined with the difficulty of linking individual crowns in remote sensing images to field data, especially for small crowns harder to detect with airborne technology (Jakubowski et al., 2013). However, pixel or plot-based output results in a disconnect between the remote sensing analysis and one of the fundamental biological units: the individual (Liu et al., 2016, De Angelis, 2018, Marconi et al. 2019). Individual plants reproduce, interact with their neighbors, and exhibit plastic responses to environmental conditions at the individual scale. Populations of individuals evolve in response to natural selection. As a result, our understanding of many biological processes is grounded in the individual and many field-based survey methods focus on collecting data with individual trees as the primary unit of measurement. While forest inventories hold information of the individual trees ineach plot (Newnham et al., 2015), plot level estimates from remote sensing typically do not include information about the relative distribution of individuals and their traits within the plot, thus reducing the amount of information about community structure. To fully understand how traits vary across space and time, and are determined by biological processes, it is important to develop approaches linking these fundamental characteristics to individual trees in ways that can be applied at scales of hundreds of km^2^.

Despite its importance for biological research, few studies have attempted to model both leaf and structural traits using remote sensing at the individual level over landscape scales (but see Chadwick & Asner 2016, Martin et al., 2018). Even when crown level models are developed, the resulting leaf trait predictions are made for pixels, not crowns, due to the challenges in crown segmentation, poor spatial resolution of hyperspectral data, or misalignment between LiDAR and hyperspectral data (Blaschke, 2010). Similarly, there are few studies estimating structural traits (like crown height and area) at crown level, with traditional methods predicting tree height and cover at the plot level (Kaartinen et al., 2012). Perhaps as a result of these differences, structural and chemical leaf traits are not currently predicted together at large scales. Consequently, ecology lacks the large scale individual level trait estimates that are necessary to fully understand tradeoffs between leaf and tree structural traits, and to explore how trait variability relates to species and the environment. Data from NEON Airborne Observatory Platform NEON (AOP) provide co-registered, georeferenced, and atmospherically corrected high resolution hyperspectral data and LiDAR, whose integration represents a great opportunity to circumvent these challenges.

To address this gap, we developed a pipeline for making crown level trait predictions at scales of ∼400 km^2^ with associated uncertainties on both crown segmentation and trait estimation. Building on Chadwick & Asner (2016) and Martin et al. (2018), we: (1) identify individual crowns in remote sensing imagery that are associated with field-based trait measurements; (2) build models relating the remote sensing data to the field-based trait measurements; and (3) apply those models to estimate trait values and examine patterns of tree structural and chemical traits from individual to landscape scales. We advance the state of the art (Chadwick & Asner 2016, Martin et al. 2018) in this pipeline by using crown-level models and comparing them to pixel- and crown average-level models, directly estimating uncertainty in trait predictions using likelihoods, and predicting traits at the crown-level. Finally, derived data products on the location, size, shape, and leaf traits of millions of individual trees distributed over tens of thousands of hectares.

## 2. Methods

In our pipeline for predicting crown level leaf and structural traits from remote sensing we used: 1) field measurements of traits for building and evaluating models; 2) data on the shape and location of individual tree crowns (ITCs) for building accurate models and assessing uncertainty in crown segmentation algorithms; and 3) high resolution remote sensing LiDAR (for crown segmentation and estimation of structural traits) and hyperspectral data (for estimation of leaf chemical traits). To obtain these components, we combined National Ecological Observatory Network’s (NEON) airborne observatory data with field data we collected at NEON sites on leaf traits as well the location and shape of individual tree crowns.

### 2.1. Site descriptions

The study was conducted at two core terrestrial NEON sites; Ordway Swisher Biological Station in Florida (OSBS, NEON Domain 03) and the Oakmulgee Management District of Talladega National Forest in Alabama (TALL, NEON Doman 08). The two sites (Appendix S1: Figure S.1) have a mix of deciduous, evergreen, and mixed forest types (Homer et al., 2012). Upland areas at both sites are dominated by fire-tolerant oaks and pine species, primarily longleaf pine (*Pinus palustris*). The longleaf pine at OSBS forms open stands whereas the longleaf pine canopy at TALL is more closed. Lowland areas near lakes or wetlands (OSBS), and riparian areas (TALL) are dominated by closed canopy hardwood forests (<u>Beckett and Golden 1982</u>, <u>Cox and Hart 2015</u>).

### 2.2. Remote sensing data

All aerial remote sensing data products were provided by the NEON Airborne Observation Platform (NEON-AOP, Table 1). We used data from the May 2014 flight for OSBS, and the June 2015 flight for TALL. We used the raw L1 data products: (1) “classified LiDAR point cloud”, and (2) “hyperspectral surface reflectance” data, orthorectified and atmospherically corrected (details in <u>data.neonscience.org/api/v0/documents/NEON.DOC.001288vA</u>). To reduce the effects of non-lambertian diffuse scattering, we applied the topographic and bidirectional reflectance distribution function (BRDF) corrections by adapting scripts from the HyTools repository (https://github.com/EnSpec/HyTools-sandbox) to our dataset. The LiDAR data consist of 3D spatial point coordinates (4-6 points/m^2^) which provides high resolution information about crown shape and height. These data are released in 1 km x 1 km tiles. Hyperspectral reflectance data consist of 1m^2^ spatial resolution images with 426 channels (or bands), each one collecting the magnitude of reflectance in 5 nm wide interval of wavelengths, ranging from visible to near infrared light (from 350 to 2500 nm). These images were provided as multiple ∼15 km x 0.8 km flight lines with a total area of ∼215 km^2^ in OSBS, and ∼145 km^2^ in TALL. The hyperspectral images were provided as “prototype” data, pre-processed differently than post 2017 data, and delivered on hard drives. Prototype airborne data showed misalignments between LiDAR and hyperspectral products, (as well as across hyperspectral flightpaths), on the scale of 1-2 meters (Marconi et al. 2019, Appendix S1: Figure S.6), primarily affecting pixels at crown borders. Despite the prototype data being potentially of lower quality than the newer NEON AOP data, we used it to match the collection dates of the field data. The only difference with current L1 and L3 data is in the nomenclature of the .h5 data structure, making the methods presented here suitable with more recent NEON data.

**Table 1.** Data products and sources (National Ecological Observatory Network, 2016). Information about data products can be found on the NEON data products catalogue (http://data.neonscience.org/data-product-catalog).

| Name | NEON data<br>product ID | Data date | How it was used |
| --- | --- | --- | --- |
| Spectrometer orthorectified at-sensor radiance | NEON.DP1.30008 | 2014 and 2015 | Hyperspectral images used to model foliar chemical and physical properties |
| Discrete return LiDAR point cloud | NEON.DP1.30003.001 | 2014 and 2015 | Crown segmentation and calculation of tree height |
| AOP L2 and L3 data products (Albedo, Elevation, Slope, Aspect) | NEON.DP2.30012.001, DP3.30024.001, DP3.30025.001 | 2014 and 2015 | Link modeled tree crown properties with other NEON-AOP data products |
| Woody plant vegetation structure | DP1.10098.001 | 2014-2018 | Build allometric relationship to infer DBH from Crown Area and Tree Height |
| Field ITC | Graves et al. 2018 | 2017 | Validate crown segmentation, define tree objects. |

### 2.3. Field Data

During this project, leaf trait data collected by the NEON Terrestrial Observation System (TOS) were not available. Instead, we used a dataset of leaf samples collected for 157 trees, many of which were near NEON inventory plots that are randomly located across the study site and stratified by land cover type. The sampled trees were located away from major roads, had crowns visible from airborne aircraft and identifiable in the image, and had sunlit branches that were accessible for leaf collection. Trees were not sampled within NEON plots to avoid disturbing NEON’s long-term monitoring efforts. As wide a range of species as possible were collected, including some rarer species that occured far from NEON plots (Appendix S1: Figure S.2).

Trait data were collected in the early part of the peak growing season in 2015. Specifically, 81 individual trees (of 17 species) were sampled from OSBS in May-June 2015, and 78 individuals (26 species) in July 2015 from TALL. Leaves were sampled from the sunlit portion of the canopy with a shotgun. Immediately after collection, the leaves were placed in a labeled plastic bag and stored in a cooler until they could be processed in the field lab within 2-4 hours of collection. The collected leaves were randomly sampled for further processing in two ways. First, a sample of leaves (at least 20 grams of fresh leaves) was analyzed for nitrogen (%N), carbon (%C), phosphorus (%P) by weight, according to standard protocols, with the exception that petioles were removed (Murphy and Riley 1962; Cornelissen et al. 2003). Second, a sample was processed for LMA using the Carnegie Institute for Science spectronomics protocol (https://gao.asu.edu/spectranomics, Asner et al., 2011). Whole leaves were weighed then scanned on a flatbed scanner to determine leaf area. The leaves were then dried at 60 C for at least 72 hours and reweighed to get the dry leaf mass. For needle-leaf species, a sample of individual needles (at least 3 fascicles per sample) was scanned and weighed. The needled dimensions of a subset of the samples were also measured with calipers to calculate total surface area. These measurements showed good agreement with the projected surface area from the scans (R^2^ > 0.75). This data and complete metadata will be added to TRY database v.6 (Kattge et al., 2020).

Individual trees were mapped in the remote sensing images using a field tablet and GIS software. Mapping was done on 2014 imagery for OSBS, and on 2015 imagery for TALL. This process involved mapping individual tree crowns on the hyperspectral image in the field to ensure the sampled trees matched directly with image pixels (Graves et al., 2018). This individual tree crown (ITC) data provides the most accurate link of field measurements with pixels from remote sensing spectral data and was used to quantify uncertainty in crown segmentation algorithms.

### 2.4. The algorithm pipeline

We developed a modular pipeline based on three steps: (1) build and evaluate crown segmentations from LiDAR data (section 2.6); (2) develop an ensemble of statistical models to infer leaf mass per area (LMA, g m-^2^), nitrogen (%N), phosphorus (%P), and carbon (%C) per tree from hyperspectral data (section 2.5), and models to estimate structural traits [diameter at breast height, DBH (cm), crown area, CA (m^2^) and stem height, H (m)] from LiDAR data; (3) make predictions for every individual tree crown in both NEON sites. For each crown, we also extracted values of elevation, slope and aspect provided as NEON AOP data products (Table 1), aiming to build a comprehensive dataset including topographic, leaf chemical and tree structural traits for any tree detected within the AOP footprint. We limited our analysis to individual tree crowns taller than 2 meters and wider than 1m^2^. Since the field traits dataset was for sunlit foliage, we predicted traits only from the upper portion of the canopy. The structure of the pipeline presented in this paper is summarized in Figure 1.

**Figure 1.**
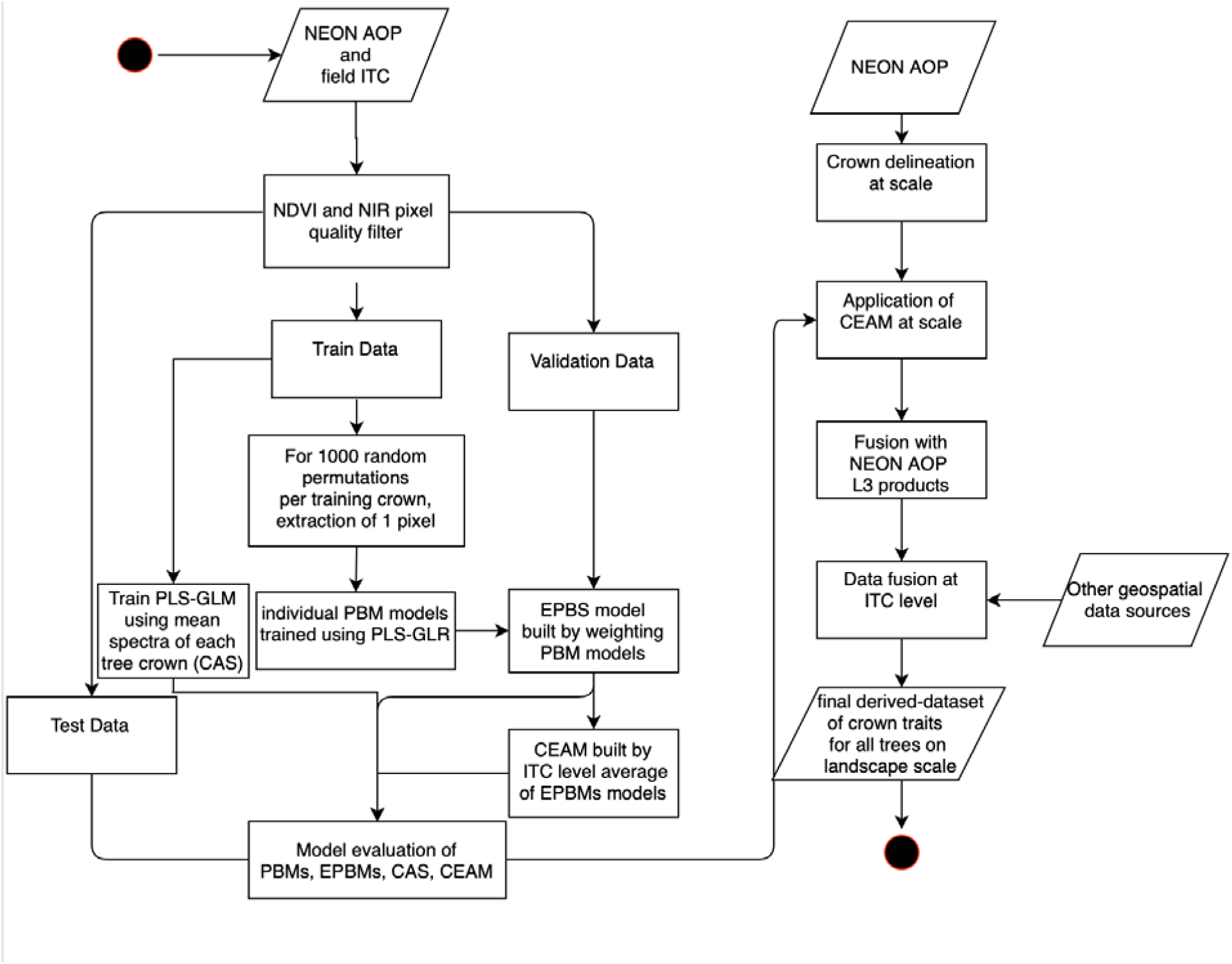
Workflow of the pipeline following Unified Modeling Language (UML). Left side shows the method to build and test three modeling approaches: PBM (pixel-based model), EPBS (ensemble of pixel-based models), CEAM (crown ensemble averaged model). Right side represents the part of the pipeline dealing with scaling, fusion, and public distribution of derived data. NEON L3 products fused to the derived traits data are: Aspect (DP3.30025.001), Elevation (DP3.30024.001), and Slope (DP3.30025.001).

### 2.5. Leaf chemistry model

After correcting the hyperspectral data with the bidirectional reflectance distribution function and topographic correction, we extracted all pixels within the boundaries of the field-delineated ITCs. Shadowed or low vegetation pixels within the ITCs were removed using thresholds for both a near infrared band (reflectance in 860 nm < 0.3) and the Normalized Difference Vegetation Index (NDVI < 0.7) (Appendix S1: Figure S.3) Colgan et al., 2012, Graves et al., 2016). We normalized the spectral values for each pixel by dividing each spectral vector by its root sum of squares. We used this method to further reduce the effect of peripheral light and shadows within each crown (Singh et al., 2015, Feilhauer et al., 2010).

Field data were split at the tree level, and stratified by species, into training (n = 115), validation (n = 18), and test sets (n = 24). Since the two sites have similar species composition, we aggregated the two datasets to build a joint model. As is common for trait studies, our field data on foliar traits was averaged to a single value for each individual tree. Most algorithms require associating a single vector of predictor variables (i.e. the spectra) to a single response value (e.g. tree crown or plot). However, individual crowns contain multiple pixels, and crowns vary in the number and quality of these pixels. In each crown, some pixels will be better for linking traits to hyperspectral signatures because they reflect light primarily from leaves, whereas other pixels include reflectance from branches, understory, or ground. To address this, we used a bagging approach (Song et al., 2013) that takes advantage of different pixel characteristics by training, weighting, and ensembling models fit to different subsets of pixels. This approach weights the predictions from models fit to different pixels to produce a more generalizable and accurate representation of the relationship between foliar traits and their spectral signatures. To capture the range of possible models from different subsets of pixels in each crown, we randomly sampled one pixel from each training crown 1000 times, and used the resulting 1000 vectors of pixels (one pixel for each crown in the dataset) to build 1000 independent partial least squares generalized linear regression (PLS-GLR) models (Bastien et al., 2005, Bertrand et al, 2014). Instead of the regular PLS regressions used in most trait modeling (e.g., Singh et al., 2015, Wang et al., 2020, Chadwick et al., 2018) we used PLS-GLR because it uses maximum likelihood estimation to calculate the regression parameters. This is an improvement over current approaches because it: (1) allows the calculation of AIC for model averaging (Burnham & Anderson 2002); (2) provides a robust measure of uncertainty in the form of a prediction interval (Christoffersen, 1998), which allows estimating the range of out of sample predictions rather than the range of mean response; and (3) does not require bootstrapping, making the method more scalable. We used a log-normal link function for all models to reflect the fact that all traits are positive numbers. We treated site (OSBS versus TALL) as a one hot encoder fixed effect (Harris & Harris, 2010) to account for site specific ancillary conditions. The number of components included in each model were determined using 5-fold cross-validation (CV) using the PRESS statistic (Tapley, 2000) on the training set. Models for each leaf trait were trained independently.

We compared four modeling strategies that varied in how the models were developed and how the models were applied for testing (Appendix S1: Table S.1). The models are labelled based on how the model was applied as “pixel based” (applied to individual pixels) or “crown based” (applied to segmented crowns). The models were built as follows: 1) a pixel-based approach with the spectra of a single pixel randomly extracted from each crown (SPM). This approach represents the case in which only the coordinates of the leaf sample are available, and spectral information can be extracted by sampling from a pixel corresponding to the stem or leaf location; A pixel based approach (hereafter referred as the “ensemble pixel based model” (or EPBM) that included information on crown identity by labelling each pixel with an individual crown identifier and using it to ensemble a selection of 100 SPM models using multi-model averaging based on delta AIC (Burnham & Anderson 2002). In this step, we selected the 100 models for each trait that best performed on the independent validation dataset (n =15). This step was fundamental to: (a) drop models that performed worse than chance (R^2^ < 0) and therefore held not meaningful relationships; (b) massively reduce the computational resources required to scale predictions to hundreds of km^2^.This approach requires crown boundaries information for training but not for application, since it applies to individual pixels; 3) a simple crown average approach, hereafter “Crown Average Spectra” (CAS), where each individual tree was represented by the average of the spectra across all green pixels (i.e. pixels with NDVI >0.7 and NIR > 0.3) within the crown polygon; 4) A crown average approach, that we refer to as the “Crown Ensemble Aggregation Model” (CEAM), consisting on averaging predictions from the EPBM for all sunlit pixels belonging to individual tree crown polygons. We tested the performance of each approach in two ways, on pixels extracted from (1) ground delineated crowns (Graves et al., 2018), and (2) algorithmically delineated crowns (Silva et al., 2016). This step was fundamental to quantify the effect of uncertainty in crown detection and segmentation on predicting leaf traits at crown level across the landscape where no field delineated crowns are available.

All models were tested on the 24 crowns withheld in the test dataset. The test data were not used at any phase of the fitting or the ensemble process. Accuracy was evaluated using the predictive coefficient of determination (R^2^) and the root-mean-square error (RMSE). The coefficient of determination produces values between 1 and negative infinity, where negative values indicate that the model predictability is lower than the sample average. As such, negative R^2^ values indicate that the statistical model did not learn any meaningful information from the data. A value of 1 indicates that predicted values perfectly match observations. We evaluated the uncertainty of predictions for each model using the coverage of the 95% prediction interval (95PI). The prediction interval is the range of values that is expected to contain 95% of the observed data points, and therefore a model with good estimates of uncertainty should have approximately 95% of the test data falling within this range. Since the CEAM was generated by the ensemble of the 100 best SPMs, we estimated the 95PI for CEAM predictions by averaging the error functions for the same 100 SPMs. We used the same data split, data transformation and PLS-GLR parameterization for all models. For pixel-based estimations (SPM and EPBM), we compared ground measures of LMA, N, C, and P with predictions from each pixel in the test dataset. For crown-based estimations (CAS and CEAM), we averaged pixel-based predictions belonging to all of the pixels in the crown. The same rationale was used for comparing pixel and crown-based uncertainty.

### 2.6. Tree structural traits and crown segmentation

We used the lidR R package (Roussel & Auty, 2017) to process point cloud LiDAR data to create a 0.5 m^2^ resolution canopy height model (CHM) and produce algorithmically delineated crowns. Despite there was little difference to the 1m^2^ resolution of NEON CHM, we chose an higher resolution CHM to produce smoother polygons and leverage the information in regions where the point cloud was more dense. We used the CHM to determine the number of trees in the scene (i.e. tree detection) using local maxima filtering (Popesco et al., 2004). We tested three alternative methods for crown segmentation (Dalponte & Coomes 2016, Silva et al., 2016, and a watershed algorithm as in Barnes et al., 2014) and chose the best performing one to generate crown boundaries (Appendix S1: Table S.2, Appendix S1: Section 1). To evaluate accuracy of crown detection and segmentation on the targeted landscapes, we calculated: (1) an estimate of precision from all predicted crowns whose boundaries overlapped with the field ITCs; (2) pairwise Jaccard index coefficient (Real & Vargas, 1996), which represents the intersection over union between the areas of two polygons, and is the standard benchmarking metric for image analysis (Rezatofighi et al., The Jaccard index was calculated by comparing ITCs collected in the field with the single most overlapping predicted crown (Marconi et al., Field delineated crowns that do not overlap with any crown segmented by the algorithm were labelled as undetected. 2019). We estimated tree structural traits from the derived polygons and the CHM. Crown area (CA) was calculated from the polygon geometry using the geoPandas python package (https://geopandas.readthedocs.io/). Tree height (H) was extracted from the CHM as the maximum height within each ITC. Diameter at breast height (DBH) was calculated using an allometric regression model relating the log-transformed DBH taken from the NEON woody plant vegetation structure data to the log-transformed height and canopy area of the matching algorithmically delineated crowns for 566 individual stems. Delineated crowns were matched to field-mapped stems in the NEON dataset visually (Appendix S1: Table S.3).

### 2.7. Building individual-level derived data for full flight paths

Each remote sensing image was split into 1 km^2^ tiles to optimize computational resources and allow parallelization on hundreds of cores. We pre-processed each tile using the same filters used for developing the models. To make predictions we used the EPBMs ensemble models to produce rasters of LMA, %C, %P and %N predictions and the 95PI for each suitable pixel and averaged them to crown level by using algorithmically delineated crowns. Crown-based predictions were achieved by averaging the values of all suitable pixels within the corresponding predicted ITC boundaries. For those areas where the ITC overlapped with more than one flight path (flight paths overlap by ∼30%), we averaged the crown-based predictions from both flight paths.

We stored flight-paths level maps of traits into raster data-products. The crown level dataset was then compiled as a comma delimited file containing all the geometry information to rebuild polygon shapes and locations. The data is distributed in a Zenodo archive (http://doi.org/10.5281/zenodo.3232978).

## 3. Results

We chose the crown segmentation algorithm described in Silva et al. (2016) (Appendix S1: Table S.2) to produce algorithmically delineated crowns. The approach detected ∼88% of the field crowns (Appendix S1: Table S.5), but showed lower accuracy in estimating the shape and size of the canopies for individual trees, with Jaccard Index ranging between 0 (for undetected trees) and 0.81, with an average of 0.35. Crowns identified by the algorithm were generally larger than those delineated in the field, resulting in overestimated crown areas (especially for smaller trees) and weak correlations between field data and algorithmic crown areas (Figure 2). Low goodness of fit in predicting Crown Area (CA) was exacerbated by uncertainty in alignment with field and remote sensing data. For example, field crowns were delineated on the hyperspectral images to incorporate only the pure pixels of the crown (Graves et al., 2018, Appendix S1: Figure S.4, Appendix S1: Figure S.5) leading to potentially underestimating the full extent of tree crown size. Moreover, visual assessment of paired field and algorithmically delineated crowns shows shifts by 1-2 meters that likely result from imperfect alignment between LiDAR and hyperspectral data (Appendix S1: Figure S.6; Marconi et al. 2019), further affecting uncertainty in field to algorithmically estimated crowns. Estimates of other structural traits from the algorithmic tree crowns were better than crown area estimates. Height showed the highest correspondence between field and remotely sensed measures (R^2^ = 0.90 for trees higher than 3m). Despite low accuracy in its predictions, CA showed a significant effect in estimating DBH from LiDAR (Appendix S1: Table S.3). However, tree height was the most important variable in predicting DBH, which therefore resulted in good estimates for both sites (0.62 R^2^), in the range of other recent applications (0.59 in Dalla Corte et al., 2020, 0.62 to 0.83 from Yao et al., 2012).

**Figure 2.**
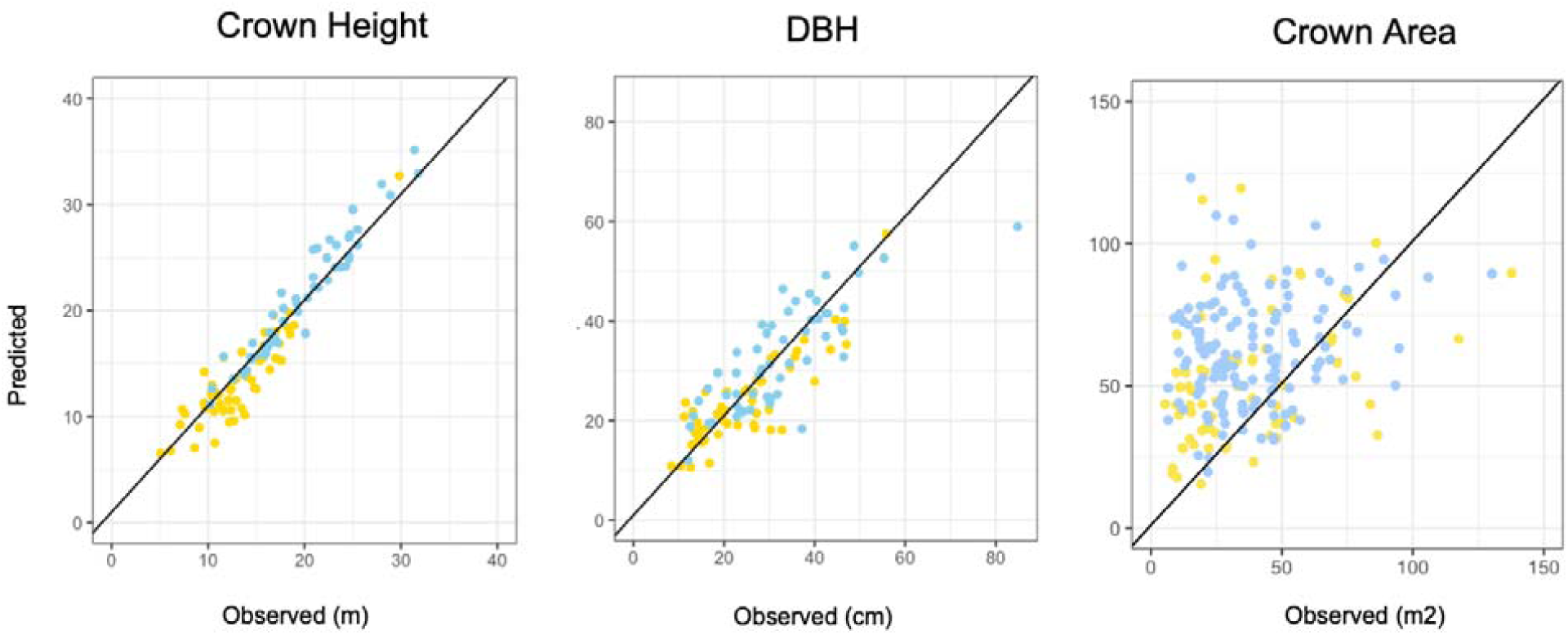
Comparison between observed and predicted tree structural traits for algorithmically delineated crowns corresponding to ground delineated ITCs: (A) tree crown height (m), (B) tree diameter at breast height (cm), and (C) tree crown area (m2). Yellow and blue points represent OSBS and TALL site data points respectively. Black diagonal is the 1:1 line.

We compared the four methods - two applied to pixels (the single pixel methods, SPM, and ensemble pixel methods, EPBM) and two applied to crowns (the crown average, CAS and the crown based ensemble methods, CEAM) - using RMSE. When tested on pixels extracted from field delineated crowns, CEAM performed the best for %N and %P (RMSE of 0.20 and 0.026), and CEAM and EPBM performed equivalently for LMA (RMSE of 40.3 and 40.8 respectively) (Figure 3A, Appendix S1: Figure S.7). CEAM explained 75% of the variance in LMA, 66% of the variance in %N, 46% of the variance in %P, and 41% of the variance in %C (Appendix S1: Table S.4, Appendix S1: Figure S.8). These results were comparable to those obtained by Martin et al. (2018) for trees in Borneo (71%, 46%, 44%, 48% respectively for LMA, %N, %P, %C). Despite “site” ancillary information was an important feature for all models, its influence on traits predictions was always relatively low compared to reflectance (as shown by the models’ parameters, Appendix S1: Figure S.9). The root mean squared error (normalized by traits observations range, NRMSE) was always between 8 and 16% of the range of the field observations, meeting the quality threshold recommended by Singh et al. (2015).

**Figure 3.**
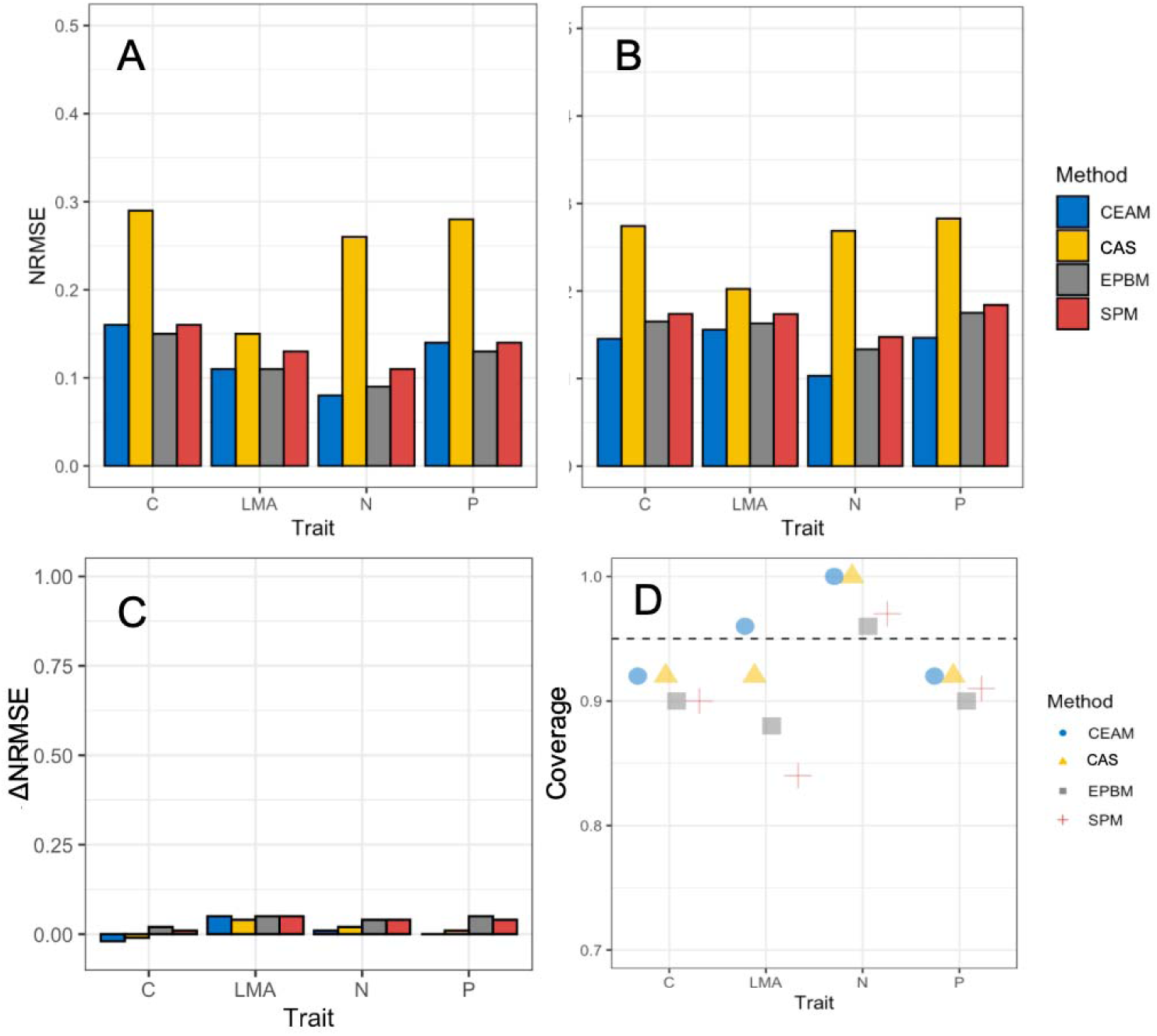
Model evaluation and comparison of pixel based (SPM, red), ensemble pixels (EPBM, grey), crown spectra average (CAS, yellow), and crown ensemble average (CEAM, blue) predictions on an independent test set of 24 observed crowns for %P, LMA, %N and %C. (A) Evaluation and comparison of RMSE from models built on pixels extracted from ground delineated crowns; (B) Evaluation and comparison of RMSE from models built on pixels extracted from algorithmically delineated crowns; (C) difference in performance (RMSE) between models tested on pixels extracted from field and algorithmically delineated crowns. Positive values mean that for that comparison model built on automatically delineated crowns performed better. (D) Coverage of the 95 predictions intervals on held out data for the four models (LMA, N, P and C). Dash-dotted line represent the ideal coverage value of 95%.

The ensemble approaches (CEAM and EPBM) performed better than methods based on pixels as the fundamental unit (SPM) or simple averaging of all green pixels in a crown (CAS) when the assessment was on field delineated crowns (Appendix S1: Table S.4; Figure 3A). Few individual SPMs performed better than the EPBM ensembles (Appendix S1: Figure S.10) and when individual pixel models did outperform the ensemble, they usually were not among the best SPM models (i.e., the models with the lowest delta AIC in the validation). This suggests that EPBM provides the best method for making out-of-sample predictions at the pixel level. CEAM generally produced the best estimates of uncertainty. CEAM 95PI showed an average coverage of 95% of held out observations, CAS 94%, EPSM 91%, Plot and SBM 90%, with the ideal value being 95% (Figure 3D).

The CEAM approach performed best when making predictions using pixels extracted from algorithmically delineated crowns (Figure 3). Compared to when crown boundaries are collected from the field, the accuracy of predictions using algorithmically delineated crowns was reduced due to the uncertainty associated with crown segmentation. However, CEAM showed the lowest reduction in accuracy compared to the other approaches (ΔNRMSE ∼ 2%, Figure 3C), resulting in the lowest NRMSE for all traits (Figure 3B).

Scaling algorithmic crown segmentation and trait estimation to the full extent of the NEON remote sensing data yielded trait predictions for approximately 5 million canopy trees for the two sites combined (Figure 4, Appendix S1: Figure S.11, S.12, S.13). Landscape patterns in traits are evident, including east-west gradients in LMA, %N, and %P at OSBS (Figure 4). At TALL, lower LMA and higher %N and %P are found in a dendritic pattern associated with the stream network (Appendix S1: Figure S.12). Some traits show a bimodal distribution at each site, which is probably related to differences in needleleaf versus broadleaf species but would need to be further tested with trait estimation coupled with species predictions. On average, OSBS showed higher %N and %P compared to TALL (Figure 5). Distributions of LMA, %N, and %P in OSBS shifted to higher values than TALL, following patterns observed from the field data (Figure 5, Appendix S1: Figure S.15). Assessing correlations between estimated structural traits, leaf traits, and abiotic environmental conditions showed strong correlations between LMA, %N, and %P, consistent with the leaf economic spectrum (Wright et al., 2004) (Appendix S1: Figure S.14). Of the environmental variables, elevation had the strongest relationship with leaf traits with leaf N and P decreasing and LMA increasing with elevation (Figures S.15, S.16). Leaf traits and tree structure were correlated at OSBS (e.g. LMA with tree H, figure 5B) but not TALL.

**Figure 4.**
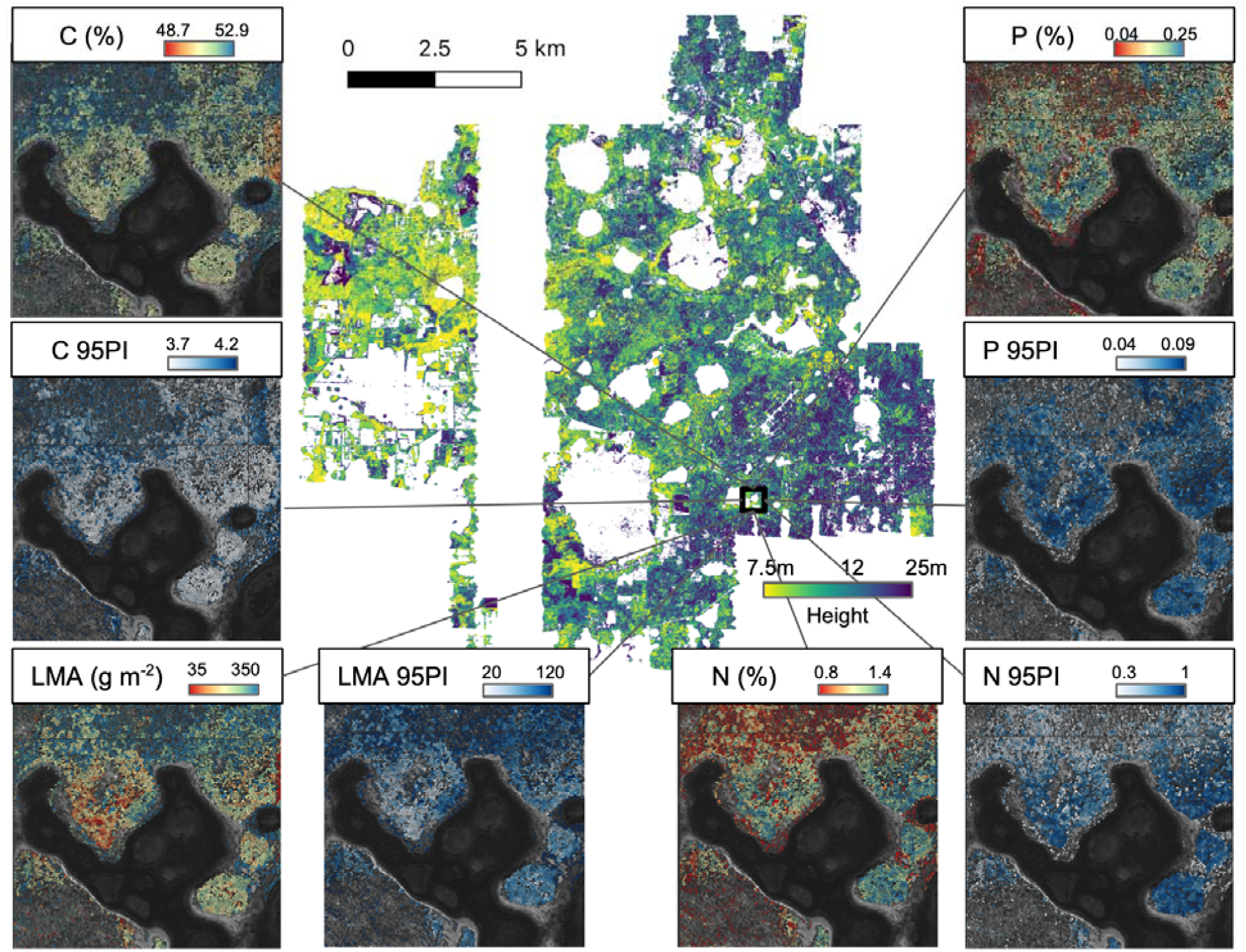
Example of predictions at the landscape scale for individual tree crowns (ITC) at Ordway Swisher Biological Station (OSBS). In the center, predictions of tree height for ∼2.5 million trees within NEON AOP footprint (215 km2), plotted in a quantile scale using the viridis color palette. Cropped images represent a ∼1 km2 detail of LMA, %N, %C, and %P predictions at scale. Expected values are presented on a quantile scale using a spectral color palette (with lower values in red, and higher values in blue). Range of the 95% probability intervals for the same area are presented on the intensity scale of blues (with lower values in white, and higher values in deep blue).

**Figure 5.**
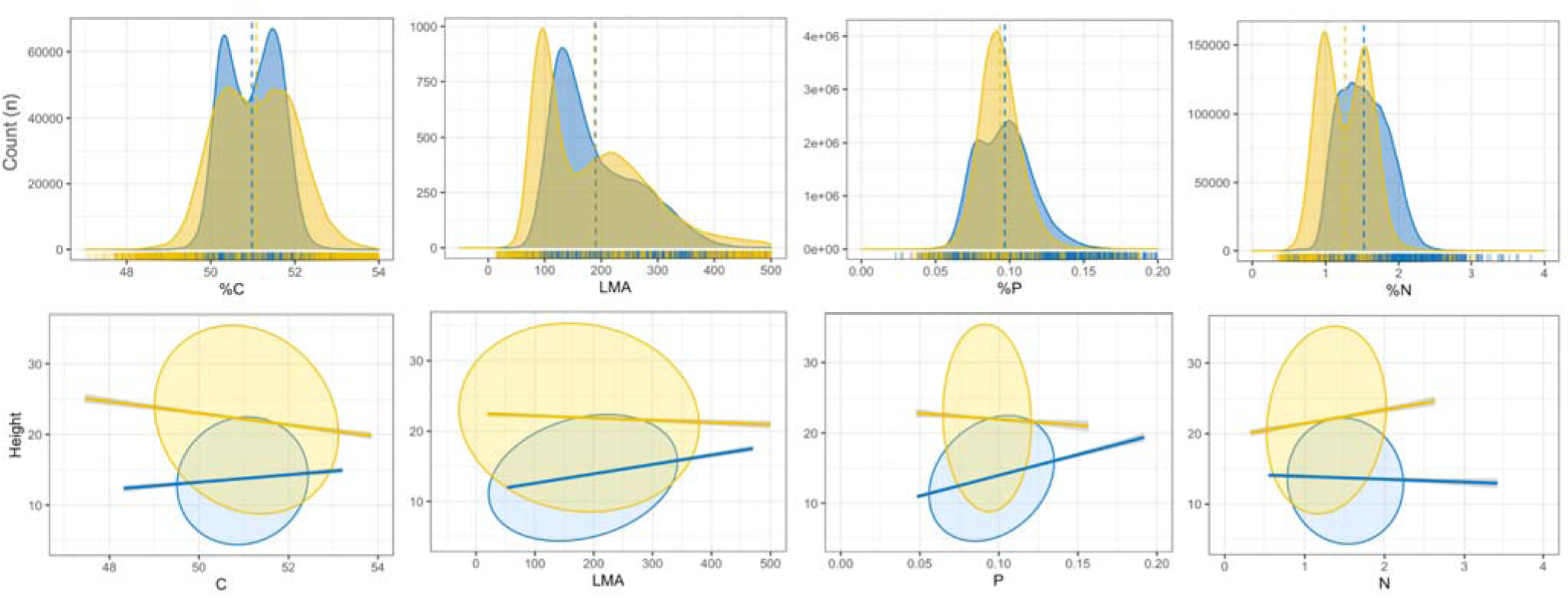
Leaf chemical distributions and relationship with tree height on a random sample of 100,000 derived individual tree crowns (ITCs). On the top row, comparison between distributions of C, LMA, %P, and %N for the two sites, OSBS (blue), and TALL (yellow). Vertical dotted lines represent the average for the site. Rug plots on the x-axis represent the marginal distribution of OSBS and TALL data between the minimum and maximum range of derived observations. On the lower row, example of relationship between tree height and the same three leaf chemical traits: from left to right, LMA, %N, %P. Linear trends and 95CI ellipses are represented for each relationship and site, following the same color scheme as above.

## 4. Discussion

The individual organism is one of the fundamental units of biology. As a result, studying the distribution of individuals and their traits across space and through time is central to many aspects of ecology. However, collecting individual level data at the large scales required for many ecological questions is challenging. To address this limitation we develop a fully automated modular pipeline to link remote sensing products from the National Ecological Observatory Network (NEON) to data collected in the field, convert the remote sensing data into estimates of the leaf and structural traits for all canopy trees detected at landscape scales, and estimate the traits for millions of individual trees in an open and accessible format (Cassey et al., 2006, Hampton et al., 2016) for use by the broader scientific community.

We found that modeling and predicting leaf traits at the individual crown level resulted in improved accuracy and uncertainty in the predictions compared to pixel based approaches (Figure 3). Linking pixels to crowns allows the ensembling of models built from the different pixels making up the crowns. Different pixels contain different combinations of leaves, branches, understory and ground, which affects the underlying chemometric relationship between foliar traits and their spectral signature. Weighted ensembling provides one way to address this, by allowing the models to identify the best combinations of pixels for relating traits and hyperspectral signatures. Aggregating pixel predictions to the crown level may also reduce the influence of outlier pixels. Our method produced models with predictive power comparable to two other crown-based estimation methods (Barbosa and Asner, 2017; Martin et al., 2018), suggesting that the performance of these approaches may generalize beyond the current study. In addition to providing robust leaf trait estimates, crown level methods allow the simultaneous estimation of structural traits, allowing these two sets of traits to be analyzed together at large scales.

2019). This is likely because height is directly measured by LIDAR and height was the most important factor in the allometric models used to predict DBH. 2018, Jucker et al. 2016). While our crown-based methods were effective for estimating a number of leaf and structural traits, there is substantial uncertainty even for the best performing traits. Quantifying this uncertainty provides information on the range of likely trait values for each individual and allows this uncertainty to be propagated when using these derived data values to test scientific hypotheses (Miller-Gulland and Shea, 2017). We used methods for leaf trait estimation that allowed us to estimate uncertainty (pls-GLR) and the crown-based method (CEAM) provided the best uncertainty estimates (Figure 3, Appendix S1: Table S.4). Current methods for delineating crowns do not include explicit measures of uncertainty (Dalponte & Coomes 2016, Silva et al., 2016). It is important for future methods to address this limitation because comparisons to field data suggested high uncertainty in segmentation.

One of the challenges for crown level approaches is that they rely on crown segmentation algorithms to identify the location and size of individual trees. While we used the best performing crown segmentation the algorithm from a recent methods competition (Marconi et al. 2019) and had reasonable correspondence between the presence of an algorithmic crown and each field crown, the algorithmic crowns averaged only 35% overlap in area with the most similar field delineated crown. Heterogeneity in point cloud density and misalignment between lidar and hyperspectral sensors could contribute to misalignment between field and algorithmically delineated crowns (Marconi et al, 2019, Kamoske et al., 2019, Appendix S1: Section 1). Despite this uncertainty, estimates of DBH and height were well correlated with field values.Therefore data derived from our pipeline should be useful for assessing estimates of individual tree biomass by applying allometric functions linking biomass to DBH and height (Graves et al. Crown area was more susceptible to the segmentation uncertainty, but is also more sensitive to small errors in segmentation and CHM resolution (Appendix S1: Section 1).

Uncertainty in crown segmentation can also have cascading impacts on the estimation of leaf traits, which was tested by comparing results from field versus algorithmically delineated tree crowns. Prediction accuracy is generally lower when using algorithmically delineated crowns because the pixels used for making predictions both include pixels that are not in the true crown and exclude pixels that are in the true crown. However, these decreases in accuracy were generally quite small, with decreases in NRMSE of <0.05 across all methods. The crown-based ensemble method (CEAM) was particularly robust to this uncertainty, with the smallest increases in NRMSE and all traits maintaining NRMSE below Singh et al. (2015)’s threshold (Figure 3C, Appendix S1: Table S.4). This robustness may result because the weighted ensembles in CEAM provide the ability to weight pixels algorithmically, allowing it to ignore pixels from outside of the true crown.

Generating derived individual level data on leaf and structural traits at the landscape scale allows trait patterns at these scales to be effectively assessed. While a complete analysis of the spatial distribution of tree traits is beyond the scope of this paper, our results showed some general patterns worth future investigations. LMA, %N, and %C showed bimodal distributions at both sites (with %N peaks particularly close in OSBS), likely because pines and oaks, the most common needleaf and broadleaf genera respectively at these two sites, occupy distinct regions of the worldwide Leaf Economic Spectrum (LES)(Wright et al., 2004). Correlation patterns between LMA and %N, %N and %P, and LMA and %P match the global scale patterns observed globally in the LES. Despite the limited number of species and geographical extent, both sites showed rangeand spread of LMA, %N and %P overlapping with most of the global range of the worldwide LES tradeoffs (Appendix S1: Figure S.14). This suggests that variation in the local environment could be driving large intra-species variability of leaf traits, while conserving the general trade-offs observed across species (Asner et al., 2016). Among the environmental variables we tested, elevation showed the strongest correlation with leaf traits (Appendix S1: Figure S.16, Appendix S1: Figure S. 17), possibly because elevation represents a proxy of different soil conditions in these sites, which can affect both species distributions and leaf traits (Walter & Gerlach, 2013). For example, small differences in elevation at OSBS often means transitioning from drained sandhill (that favor pines) to marshy and richer soils that favor the establishment of large-crowned broadleaf species, rich in foliar %N and %P (Bodker et al., 2015).

Producing derived data at the individual level also facilitates landscape scale assessments of relationships between leaf and structural traits at the level of individual organisms. For example, although the two sites have similar species composition, our results showed different correlational patterns between structural (height and DBH) and chemical traits (LMA, %N, %P, %C) (Figure 5, Appendix S1: Figure S.17), especially the relationship between height and LMA (r = 0.24 in OSBS, r = -0.08 in TALL). Possible explanations for these relationships may be related to differences in management histories across patches of the landscape that can influence species assembly, successional stages, which are important determinants of tree size and leaf traits (Sameulson and Stokes 2012; Ishida et al. 2005). Our pipeline, integrated with further remote sensing derived information (e.g. species identities) and local history (e.g. management and fire history) could be used to address how these drivers affect local distribution of plant traits, their trade-offs, and their effects on the ecosystems across a multitude of landscapes.

Our crown-based approach to modeling and predicting tree traits produces individual data similar to that collected in the field. This approach has a number of benefits. First, it will make data integration with field-based forest and trait surveys easier because both derived remote sensing data and field surveys will be composed of the same fundamental unit (individual trees). Second, crown-based approaches are likely better for aligning trait data across years. The same pixel in two consecutive years could vary significantly in a trait because of small errors in spatial alignment of pixels through time, whereas large crown-level regions will be more robust to small errors at the edges of the crown. Moreover, multi-temporal and multi-sensor images can be potentially leveraged to align and improve segmentation for crown objects captured in the same scene (Bovolo & Buzzone, 2017, Sumbul et al., 2020). Finally, this approach allows a more compact representation of derived trait data in tabular formas spatial polygons instead of rasters.

While this will not be the best representation for all analyses, for individual level analyses it results in vastly reduced storage computational requirements compared to raster data.

Thanks to the modular nature of our approach, crown segmentation can easily be substituted with methods based on RGB (Weinstein et al., 2020) or hyperspectral imaging (Dalponte et al., 2016) when LiDAR data is poor or not available. Yet, individual level approaches are limited by data availability and are not suitable for addressing ecological questions at continental to global scale. High resolution airborne remote sensing is still limited to relatively few sites, while the resolution of AVIRIS or satellite data is more appropriate for plot level analyses (e.g., Singh et al. 2015, Martin et al. 2018, Ma et al. 2019). However, these two approaches can be potentially integrated to scale sub-pixel properties from satellite data, and merge the gap between local, regional and global scale ecological information, and better address emergent cross-scale ecological questions related to variation of leaf traits, diversity and functions (Carmona et al., 2016).

The data produced by our individual level pipeline could be extended by including predictions for species identity, other leaf and structural traits, environmental variables, management, or disturbance. Moreover, our pipeline can potentially be used to extract ecological information for every tree that can be detected across all NEON AOP sites for the full life of the observatory. This will produce a publicly available, spatially explicit database of detailed ecological information for hundreds of millions of trees across the US that could be fused with other continental data (e.g. Forest Inventory and Analysis) and integrated to area based analyses from satellite data, to address cross scale functional ecological questions (http://doi.org/10.5281/zenodo.3232978). Such data could be used to further understand the biology behind trait tradeoffs and investigate cross scale ecological processes and patterns from individual to landscape to continental scale.

## 5. Acknowledgments

This work was supported by the Gordon and Betty Moore Foundation’s Data-Driven Discovery Initiative through grant GBMF4563 to E.P. White and by the National Science Foundation through grant 1926542 to E.P. White, S.A. Bohlman, A. Zare, D.Z. Wang, and A. Singh; by the NSF Dimension of Biodiversity program grant (DEB-1442280) and USDA/NIFA McIntire-Stennis program (FLA-FOR-005470) to S. A. Bohlman; by the University of Florida Biodiversity Institute (UFBI) and Informatics Institute (UFII) Graduate Fellowship to Sergio Marconi. There was no additional external funding received for this study.

## 6. Description of author’s responsibilities

Sergio Marconi and Sarah Graves designed the experiment, Sarah Graves carried out field work, Sergio Marcon, Sarah Graves and Ethan White developed the methods, Sergio Marconi performed the analysis, Ethan White and Stephanie Bohlman supervised the work as lab leaders, helped with experimental design and advice on data analysis. Ben Weinstein helped with technical aspects and manuscript editing. All authors contributed to editing the manuscript. We acknowledge Dr. Adytia Singh for his suggestions in the initial phase of the project, and the reviewers that contributed to significantly improve this manuscript.

