## Appendix 1 for "Estimating individual level plant traits at scale"

**Appendix S1.**

Title**: Estimating individual level plant traits at scale**

Authors: Sergio Marconi^1^*, Sarah J. Graves^2,3^, Ben. G. Weinstein^4^, Stephanie Bohlman^2^, Ethan P. White^4^

Journal: **Ecological Applications**

^1^School of Natural Resources and Environment, University of Florida, Gainesville, FL, 32611 USA

^2^School of Forest Resources and Conservation, University of Florida, Gainesville, FL 32603, USA

^3^University of Wisconsin-Madison, Nelson Institute for Environmental Studies, Madison, WI, United States, 53706

^4^Department of Wildlife Ecology and Conservation, University of Florida, Gainesville, FL 32603, USA

**Section S1: Evaluation of crown segmentation.**

We applied three unsupervised crown segmentation algorithms (Dalponte & Coomes 2016, Silva et al., 2016, and a watershed algorithm as in Barnes et al., 2014) to generate crown boundaries**.** We used a single parametrization (Table S1) for each method at both sites to facilitate cross-model and cross-sites comparison. Data were processed using the lidR R package (Roussel & Auty, 2017). To address the accuracy in detecting tree crowns we calculated the (1) recall and precision using 3 thresholds (10, 20, 30%) to estimate the ability in detecting trees; (2) the Jaccard index (i.e. intersection over union between field and algorithmic crowns) to estimate the ability in estimating crowns shape and area. The Jaccard index was calculated by comparing ITCs collected in the field with the single most overlapping predicted crown. Field delineated crowns that do not overlap with any crown segmented by the algorithms for more than 10% of their surface were labelled as undetected. The Silva et al. (2016) segmentation algorithm performed the best on our data (Table S1, see results section), consistent with results from a multi-group data science competition (Marconi et al. 2019). Therefore, this algorithm was used to extract ITCs from the CHM data for the full remote sensing footprint for both NEON sites.

The crown segmentation algorithm described in Silva et al. (2016) yielded the highest overlap with the field-delineated ITC crowns (Table S.2). The approach detected 88% of the ground delineated crowns (using a 10% threshold of overlapping). For individual trees, the Jaccard Index ranged between 0 (undetected trees) and 0.81, with an average of 0.35 and median of 0.34.

Low goodness of fit in predicting Crown Area (CA) was exacerbated by uncertainty in alignment with field and remote sensing data. For example, field crowns were delineated on the hyperspectral images to incorporate only the pure pixels of the crown (Graves et al., 2018), leading to potentially underestimating the full extent of tree crown size. Moreover, visual assessment of paired field and algorithmically delineated crowns shows shifts by a couple of meters that are likely a result of imperfect alignments between LiDAR and hyperspectral data (Figure S.6).

While this level of overlap suggests that improved methods for crown delineation are needed, it is also the result of limitations in the resolution and alignment of the remote sensing products. The resolution of the remote sensing data (1m2 pixels) relative to the scale of most tree crowns (average tree crown area in this study = 35 m2) means that even single pixel errors can have a large influence. For example, if the extent of a 4x4 m crown (area = 16 m2) is overestimated by a single pixel in each direction (6x6m = 36 m2), the Jaccard index is only 0.44 and CA more than doubled. In addition, remote sensing data from multiple sensors often have imperfect alignments between products. The prototype NEON AOP data used in this study may have misalignments between LiDAR and Hyperspectral products on the scale of 1-2 meters (Marconi et al. 2019). Misalignment of this extent will primarily affect pixels at crown borders. When estimating plant traits, the key question is how the errors in delineation driven by models, resolution, and alignment influence the accuracy of structural and leaf trait models. While crown-based leaf trait models are robust to these errors (see above) the structural trait estimates are directly related to crown delineation and are therefore strongly influenced by errors in that delineation. Fortunately estimates of DBH and height were well correlated with field values.

**Table S1**

| Approach | Description | Requires ITCs for training | Requires ITCs for making predictions | Tested on |
| --- | --- | --- | --- | --- |
| PBM | Random extraction of one pixel per crown to build chemometric models. | NO | NO | Pixels (n = 750) |
| EPBM | Ensemble of a number of PBMs to create a robust multiple instance approach | YES | NO | Pixels (n = 750) |
| CAV | Average of the spectra of all green pixels in a crown | YES | YES | Crowns (n = 26) |
| CEAM | Average of the PBM predictions on all green pixels in a crown | YES | YES | Crowns (n = 26) |

*Table S.1. Schematic description of the four approaches used compared in the study*

**Table S2**

| Min. | 25th percentile | Median | Mean | 75th Percentile | Max |
| --- | --- | --- | --- | --- | --- |
| Barnes et al., 2014 | | | | | |
| 0 | 0.2 | 0.3 | 0.31 | 0.43 | 0.72 |
| Dalponte & Coomes 2016 | | | | | |
| 0 | 0.24 | 0.33 | 0.34 | 0.44 | 0.78 |
| Silva et al., 2016 | | | | | |
| 0 | **0.25** | **0.35** | **0.35** | **0.45** | **0.81** |

*Table S2. Comparison of the Jaccard Index measure for the overlap between the  ground-delineated ITC crowns and algorithmically delineated ITC crowns.*

**Table S3**

| (A) | Parameter | Std.Error | DF | t-value | p-value |
| --- | --- | --- | --- | --- | --- |
| (Intercept) | -4.446325 | 3.124267 | 454 | -1.423158 | 0.1554 |
| CHM | 1.645523 | 0.084261 | 454 | 19.52896 | 0 |
| CA | 0.098375 | 0.020195 | 454 | 4.871191 | 0 |
| (B) | AIC | BIC | logLik | Obs | Groups |
|  | 3127.317 | 3147.919 | -1558.659 | 458 | 2 |

*Table S.3. Parameters of the allometric relationship derived from linking dbh measured by NEON woody vegetation structure dataset (n = 458) to tree height and crown area predicted by the pipeline (R2 on 108 held out data points of 0.62).*

**Table S4**

|  | R2 | | | | RMSE | | | | Coverage | | | | %RMSE | | | |
| --- | --- | --- | --- | --- | --- | --- | --- | --- | --- | --- | --- | --- | --- | --- | --- | --- |
|  | Tested on field delineated crowns (n = 24) | | | | | | | | | | | | | | | |
|  | PBM | CAS | EPBM | CEAM | PBM | CAS | EPBM | CEAM | PBM | CAS | EPBM | CEAM | PBM | CAS | EPBM | CEAM |
| N | 0.54 | 0.53 | 0.67 | **0.72** | 0.27 | 0.63 | 0.22 | **0.2** | **0.97** | 1 | **0.96** | 1 | 0.11 | 0.26 | **0.09** | **0.08** |
| P | 0.36 | 0.38 | 0.48 | **0.48** | 0.03 | 0.05 | **0.03** | 0.03 | 0.91 | 0.92 | 0.9 | **0.92** | 0.14 | 0.28 | **0.13** | **0.14** |
| C | 0.15 | 0.39 | 0.26 | **0.41** | 1.26 | 2.27 | **1.17** | 1.22 | 0.9 | 0.92 | 0.9 | **0.92** | 0.16 | 0.29 | **0.15** | **0.16** |
| LMA | 0.68 | 0.72 | **0.75** | **0.75** | 45.56 | 54.88 | **40.33** | 40.8 | 0.84 | 0.92 | 0.88 | **0.96** | 0.13 | 0.15 | **0.11** | **0.11** |
|  | Tested on algorithmically delineated crowns (n = 24) | | | | | | | | | | | | | | | |
| N | 0.38 | 0.49 | 0.49 | **0.6** | 0.36 | 0.65 | 0.32 | 0.25 | 0.88 | 0.95 | 0.88 | 0.86 | 0.15 | 0.27 | 0.13 | **0.1** |
| P | 0.27 | 0.4 | 0.34 | **0.44** | 0.04 | 0.06 | 0.03 | 0.03 | 0.86 | 0.91 | 0.81 | **0.91** | 0.18 | 0.28 | 0.18 | **0.15** |
| C | -0.07 | 0.26 | 0.03 | **0.28** | 1.36 | 2.15 | 1.3 | 1.14 | 0.87 | 0.95 | 0.85 | **0.91** | 0.17 | 0.27 | 0.17 | **0.15** |
| LMA | 0.52 | 0.61 | 0.58 | **0.6** | 62.2 | 72.46 | 58.43 | **55.83** | 0.77 | 0.91 | 0.8 | **0.91** | 0.17 | 0.2 | 0.16 | **0.16** |

*Table S.4 Models performance on 24 held-out crowns (predicted R2 and RMSE) for the four modeling strategies CEAM (Crown ensemble average model), Crown average spectra (CAV), EPBM (Ensemble pixel based model) and SPM (single pixel model). Higher performance in bold. (a) Train, validation and test pixels extracted from ground delineated crowns (least uncertainty in pixel labeling). (b)  Train, validation and test pixels extracted from automatically delineated crowns (highest uncertainty in pixel labeling).*

**Table S5**

|  | Precision | Recall | True positive | False Negative | False positive |
| --- | --- | --- | --- | --- | --- |
| Overlapping with 10% of a field crown | | | | | |
| TALL | 0.95 | 0.82 | 72 | 4 | 16 |
| OSBS | 0.81 | 0.84 | 65 | 15 | 13 |
| FULL | 0.88 | 0.83 | 137 | 19 | 29 |
| Overlapping with 20% of a field crown | | | | | |
| TALL | 0.69 | 0.81 | 63 | 15 | 28 |
| OSBS | 0.61 | 0.74 | 60 | 21 | 38 |
| FULL | 0.65 | 0.77 | 123 | 36 | 66 |
| Overlapping with 30% of a field crown | | | | | |
| TALL | 0.65 | 0.76 | 59 | 19 | 32 |
| OSBS | 0.59 | 0.72 | 58 | 23 | 40 |
| FULL | 0.62 | 0.74 | 117 | 42 | 72 |

*Table S.5 Crown detection scores of Recall and Precision applied to algorithmically delineated crowns overlapping with field delineated crowns. Note, precision and recall are calculated on an arbitrary threshold of overlap with ground-truth object. There is no general consensus about which threshold is better fit to crown objects, and it is not always clear which threshold is used in different studies. For this reason, we are presenting here the results at three different levels of overlap: 10, 20, 30%.*

**Figure S1**

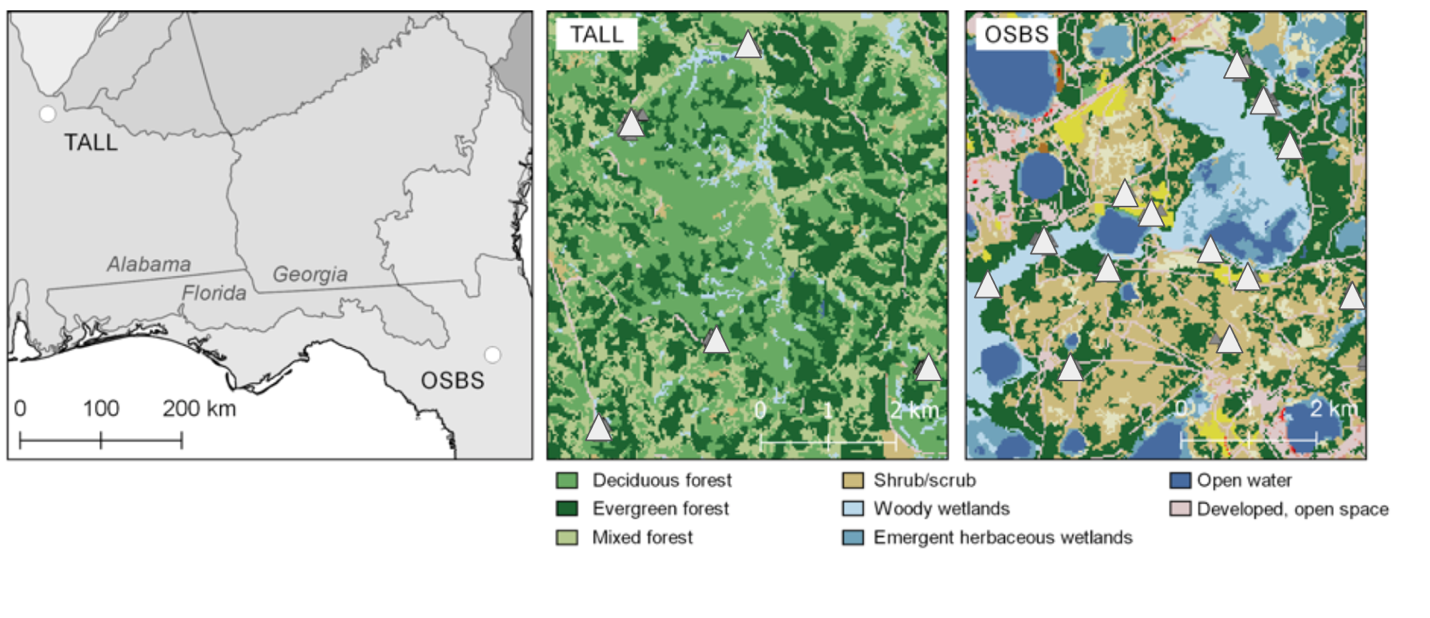

*Figure S.1. Regional map showing OSBS and TALL locations in the southeastern US ecoregions. Insets show TALL and OSBS locations of individual tree crowns (gray triangles) over the 2011 National Land Cover Database.*

**Figure S2**

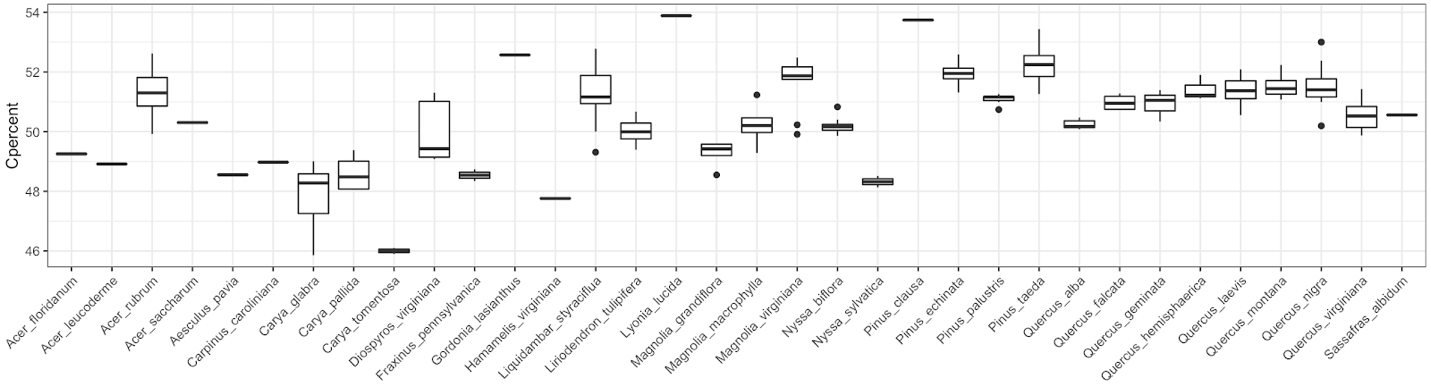

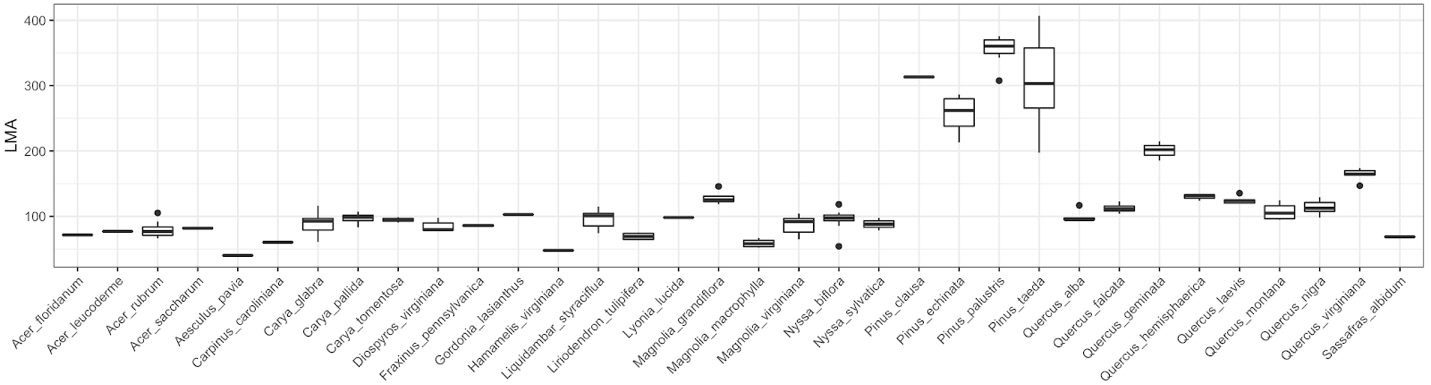

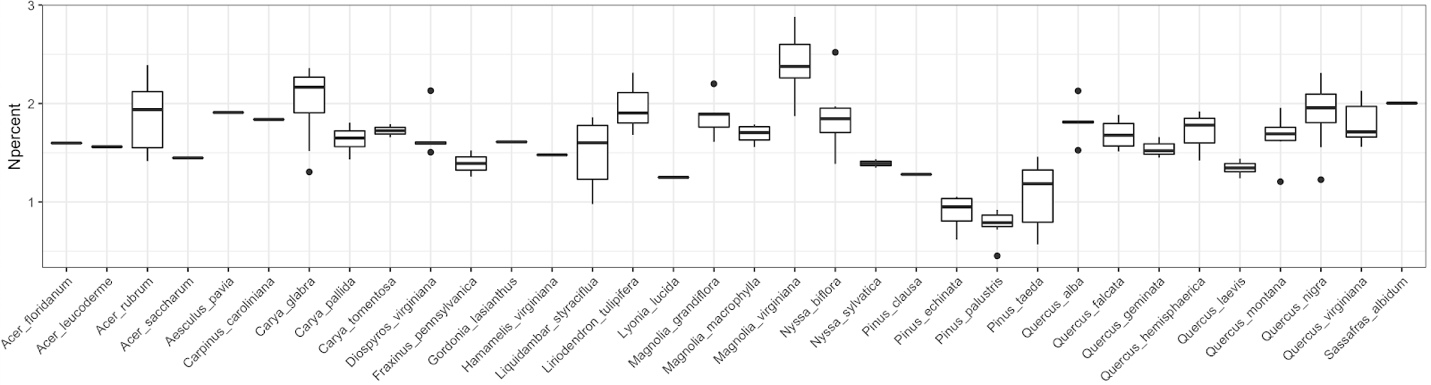

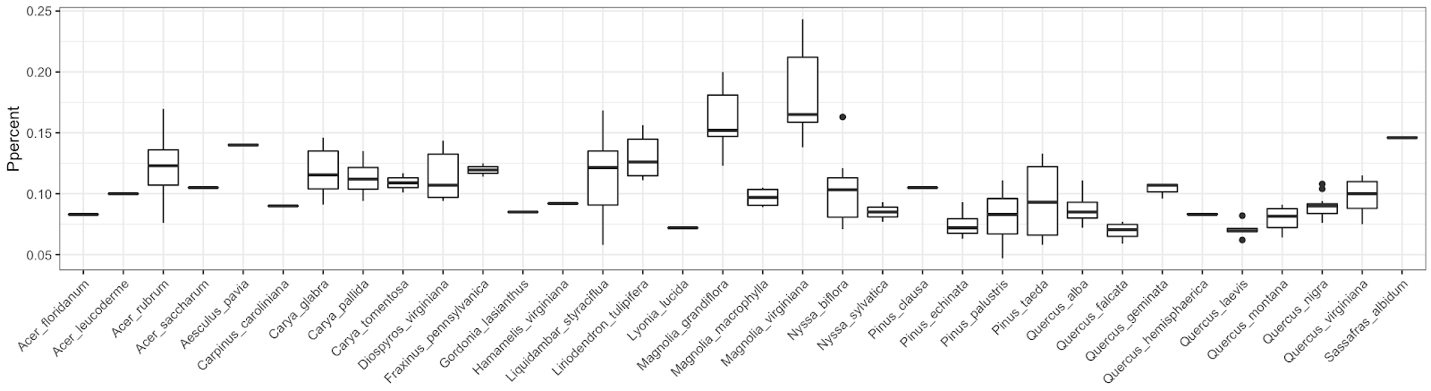

*Figure S.2 Variability in observed values in the four traits analyzed in this work (respectively LMA, %C, %N, and %P). Boxplots represent distribution of values per species sampled (34).  10 species were sampled only once, therefore could not be represented in the validation and test. Random out of sample dataset was extracted making sure that most of the remaining species were represented (22 out of 24).*

**Figure S3**

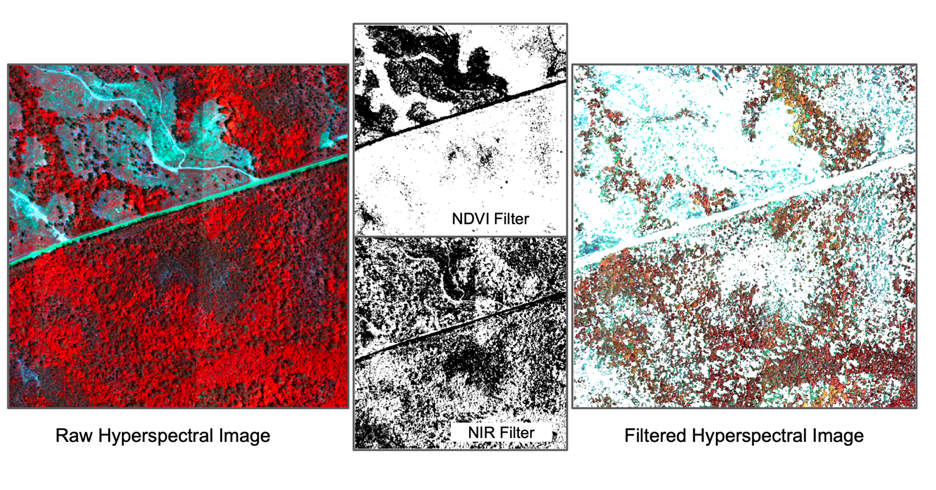

*3. False color image from hyperspectral image before and after removal of pixels that do not meet the pixel screening threshold. Left panel represents the raw hyperspectral data in false colors (bands 17, 87, 117). Center panels show the NIR (top) and NDVI (bottom) masks where white pixels were removed. Right panel shows the result of filtering. Only pixels with NDVI > 0.7 and reflectance in 860 nm > 0.3 were kept.*

**Figure S4**

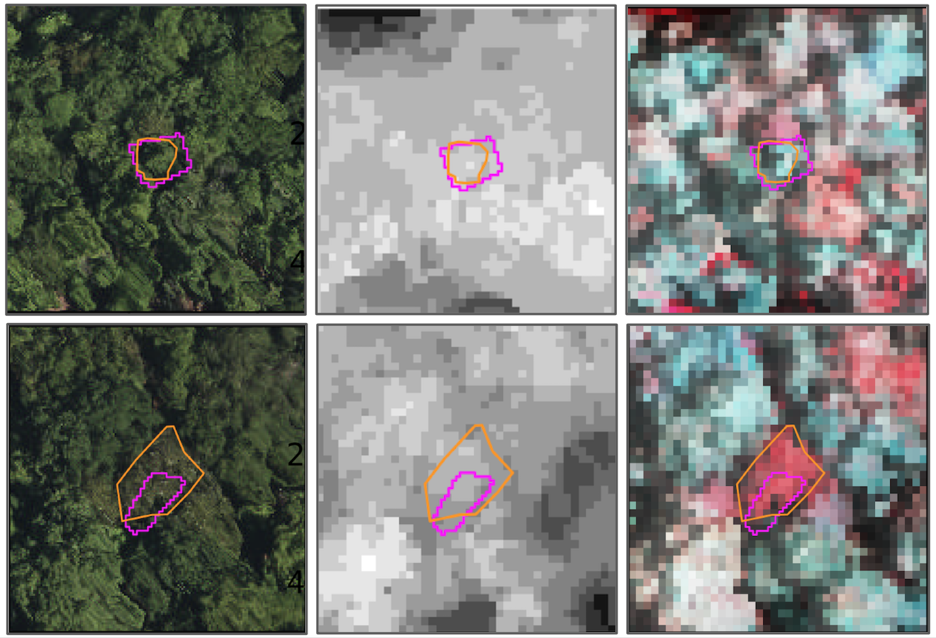

*Figure S.4. Example of relatively good (top row) and less precise (bottom row) of algorithmic delineated crowns. Yellow polygons represent ground truth data. Magenta polygons represent the single predicted ITCs overlapping with the ground truth crown. Panels represent a 40x40 m centered on the field crown centroid. Left panel represent the scene in RGB, the center panel is the NEON canopy height model, and the right panel is a false color composite RGB image (bands 17, 87, 117) superimposed on the CHM.*

**Figure S5**

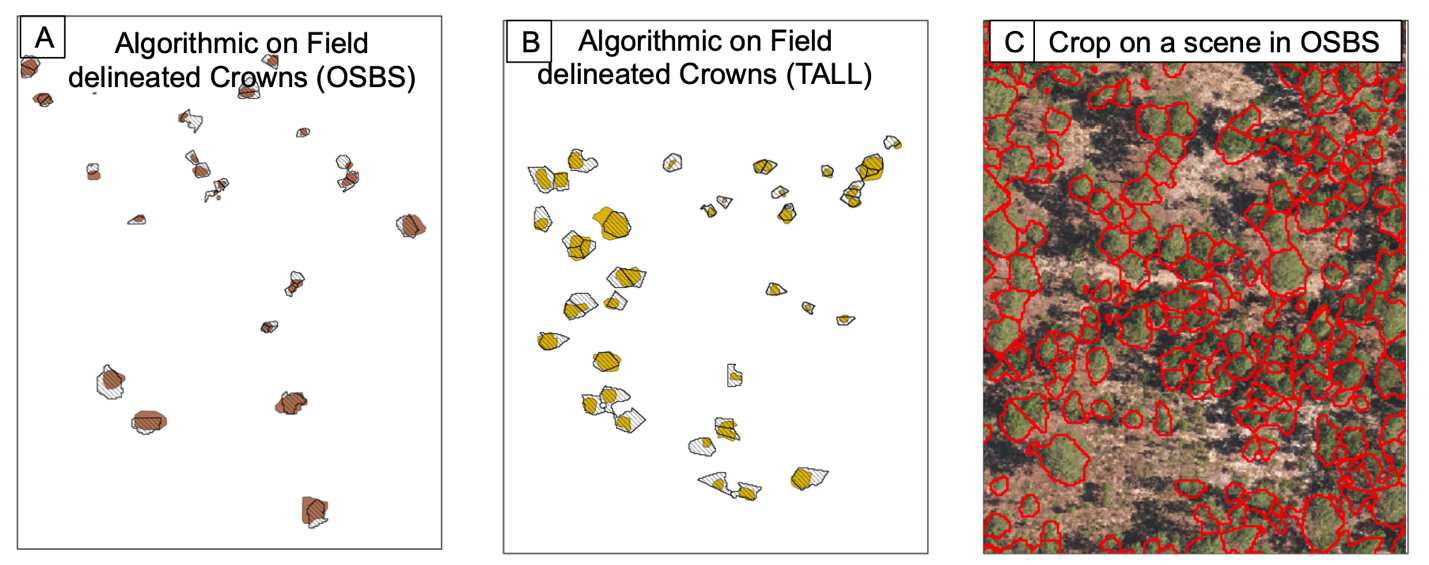

*Figure S.5. Example of segmentation results for OSBS (Box A) and TALL (Box B). Fill colored polygons represent the field delineated crowns (brown for OSBS, yellow for TALL); Hollow barred black polygons represent the algorithmically delineated crowns. Despite crowns are generally detected, algorithmically delineated crowns show over segmenting (splitting a field crown in multiple polygons), and area overestimation (especially for trees with smaller size). The left panel show an example of segmentation for a longleaf pine section in outside of the training area. Red polygons represent individual algorithmically delineated crowns. The background is a L3 Orthophoto (resolution of 0.25m) from NEON AOP data.*

**Figure S6**

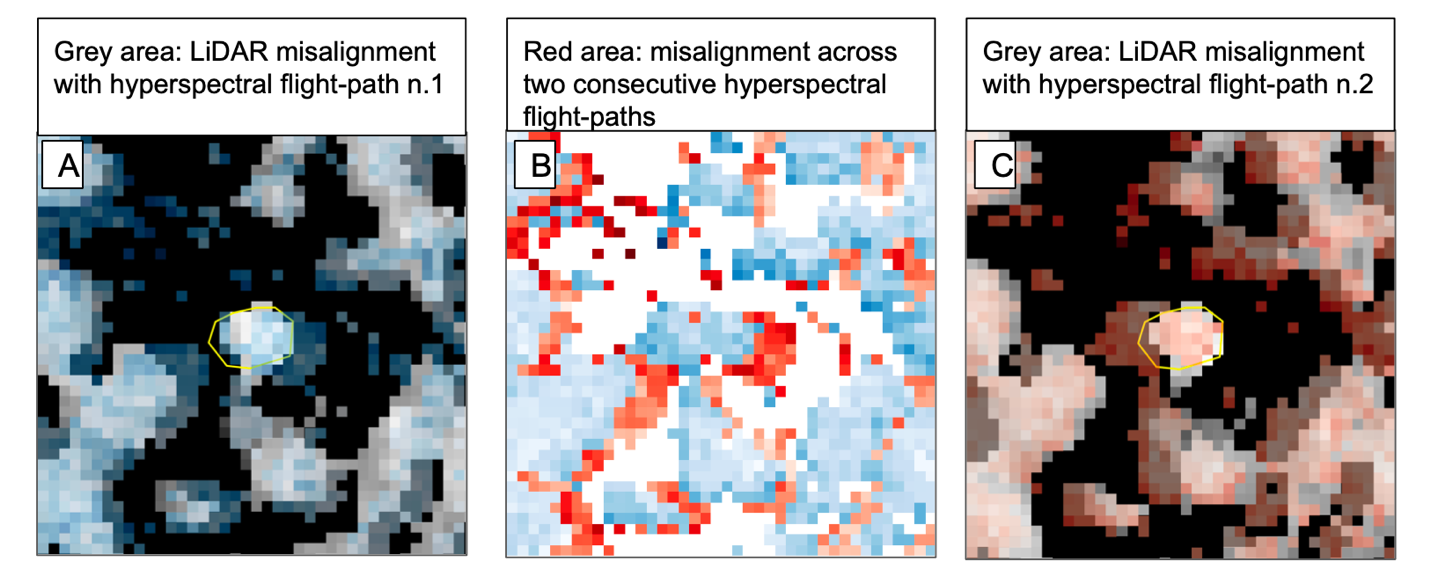

*Figure S.6 Example of misalignment between LiDAR data (here represented by a 1m^2^ CHM, in grayscale), and two different hyperspectral images registered at two different flightpaths (blue and red). (A) Misalignment between overlapping CHM and the first flightpath, showing a shift east of the hyperspectral image; (B) Misalignment between two consecutive flightpaths, showing an offset up to 4m; (C) Misalignment between overlapping CHM and second HSI flightpath, showing a shift west of the hyperspectral image.*

**Figure S7**

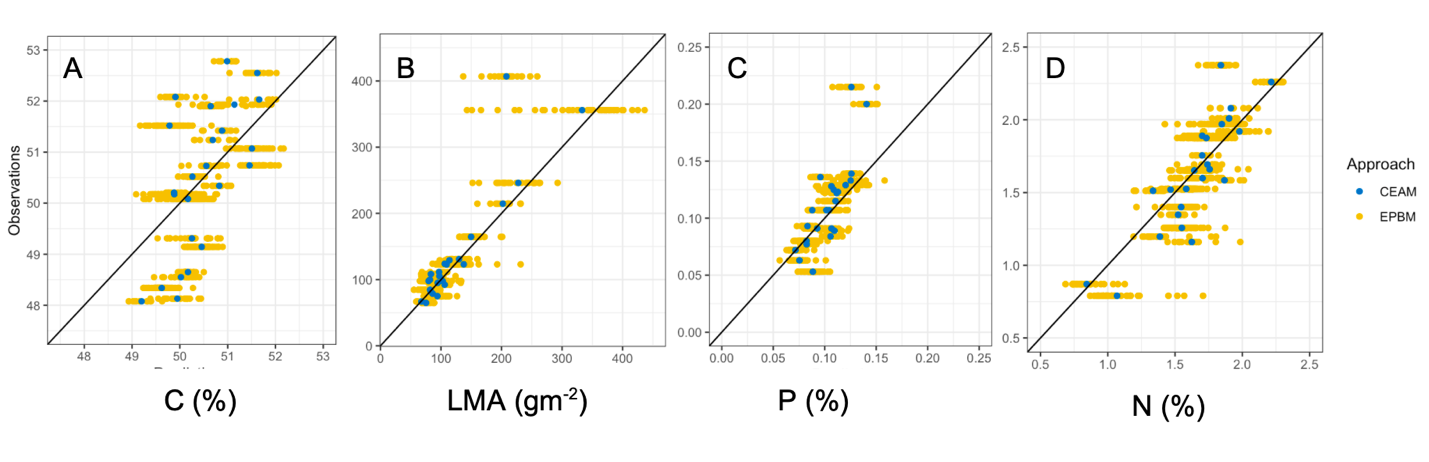
*. (A-D) Comparison of EPBM (yellow) and CEAM (blue) predictions on 24 held out observed crowns for %C (A), LMA (B), %P (C)  and %N (D) respectively. Black diagonal is the 1:1 line.*

**Figure S8**

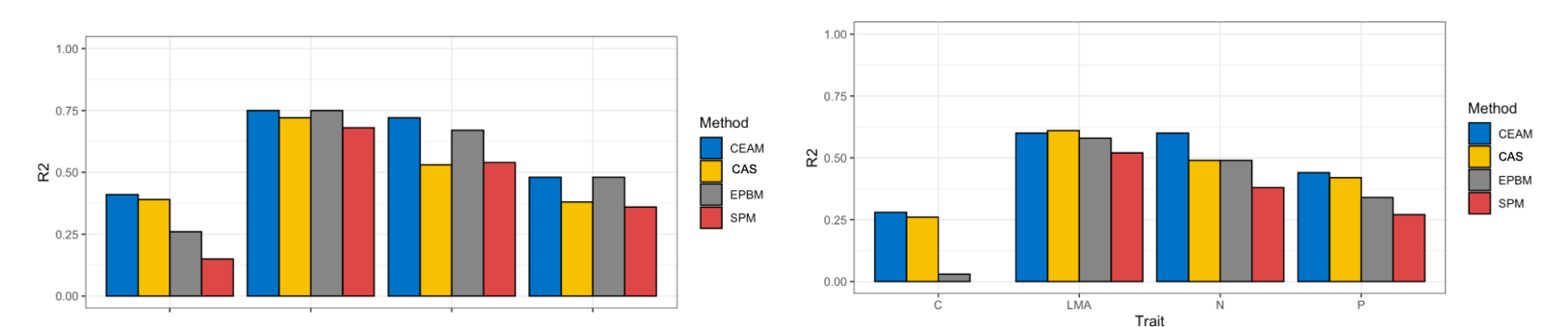

*Figure S.8. Evaluation and comparison of R2 on held out test data (24 crowns) for the four approaches: (A) models built on pixels extracted from ground delineated crowns; (B) models built on pixels extracted from algorithmically delineated crowns*

**Figure S9**

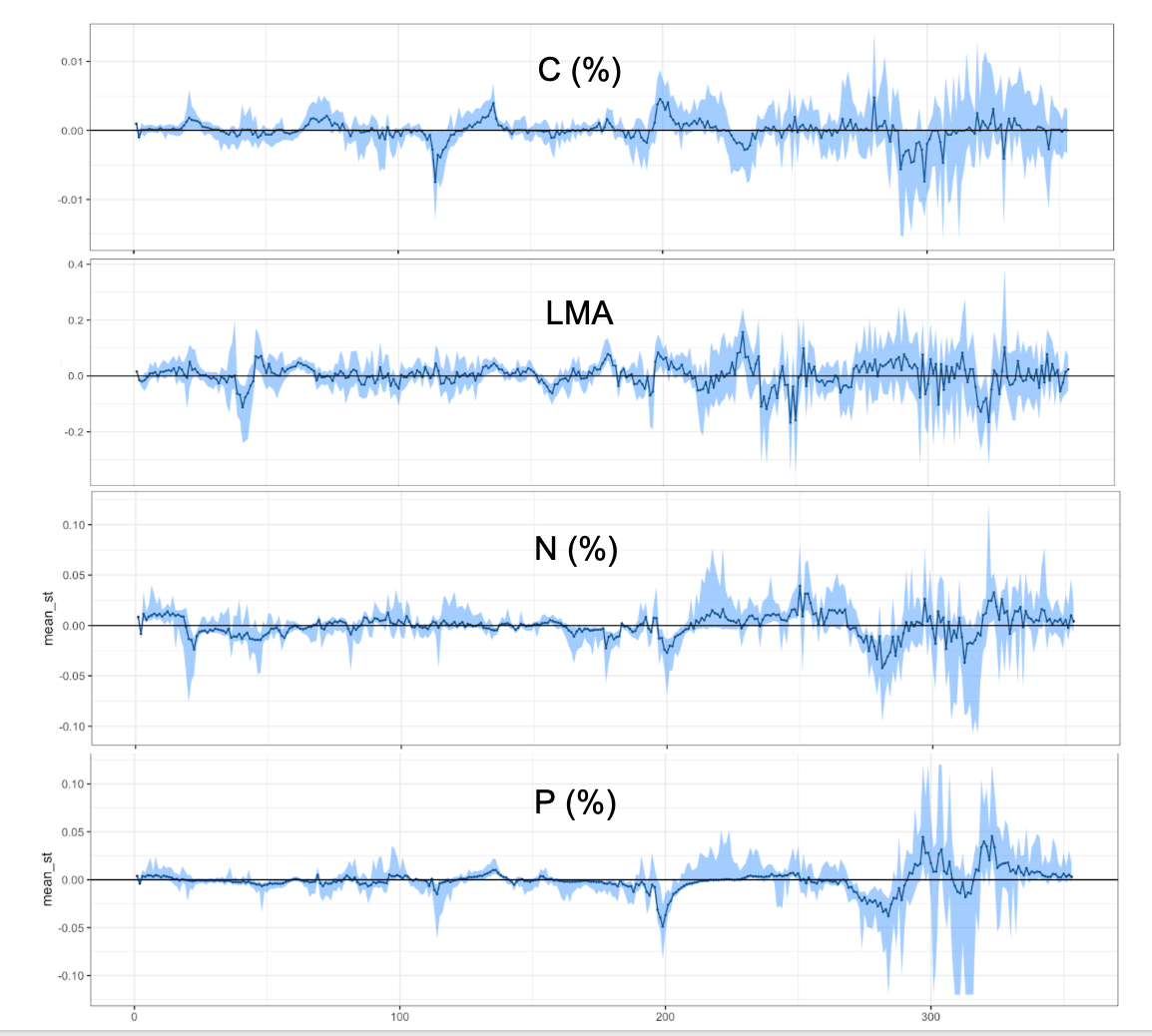

*Figure S.9 Distribution of coefficients of the 100 PBMs used to build the EPBM model. Plots show the 80% range of the values of PLS-GLM parameters for each band. Wavelengths whose interquartile range includes the value of 0 are scarcely or no informative. Note that ancillary site effects (first two bands) showed significant effect on all traits but relatively little influence compared to reflectance.*

**Figure S10**

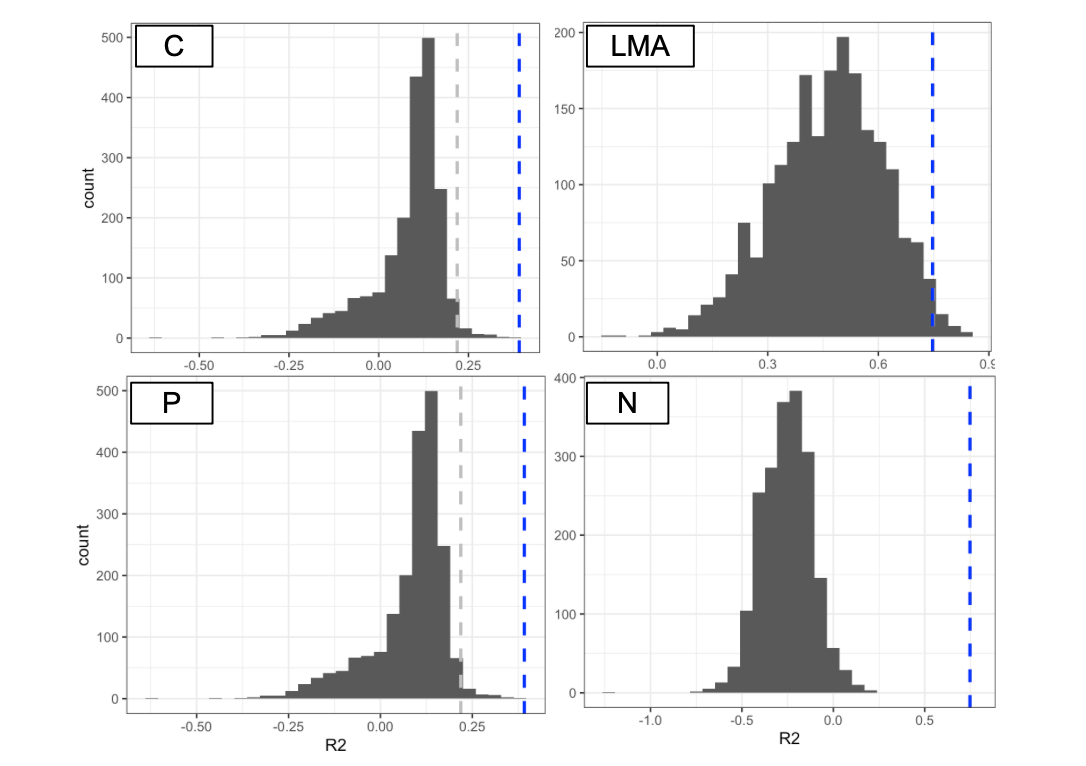

*Figure S. 10. Comparison of coefficients of determination on the validation data, using the 100 best SPMs (histogram), the EPBM (grey dashed line), and the CEAM (blue dashed lines), assessed based on field delineated crowns. An R2 of 0 represents the threshold below which the residual sum of squares (RSS) is higher than the total sum of squares (TSS). A negative R2 value means that the observed sample mean is a better predictor than the model, and so that specific model is meaningless.*

**Figure S11**

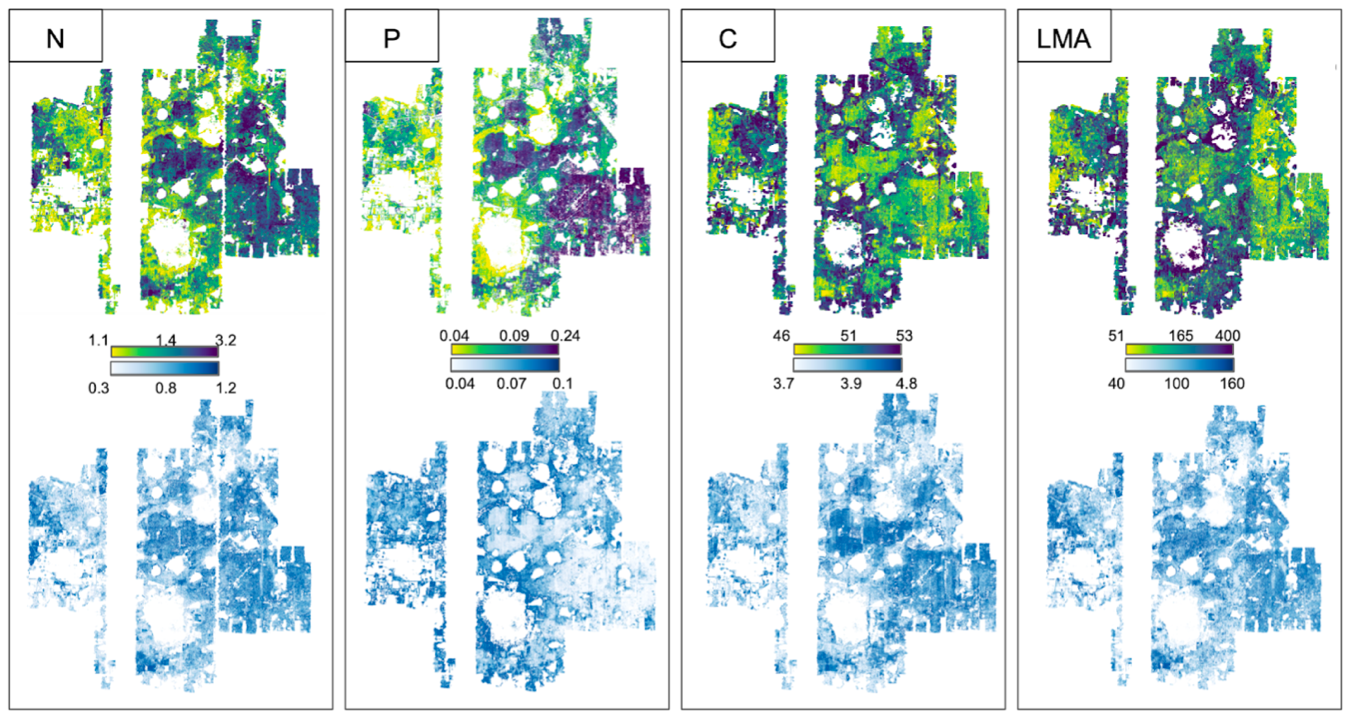

*Figure S. 11. Predictions maps for LMA, %N and %P for the OSBS site (215 km2). The maps show the centroid of around 2.5 million trees, colored by quantile gradient. The top panel (viridis palette) represents individual predictions for %N, %P, %C and LMA. The lower panel (blue) represents the uncertainty for each tree-based prediction, plotted as the width of the 95%PI.*

**Figure S12**

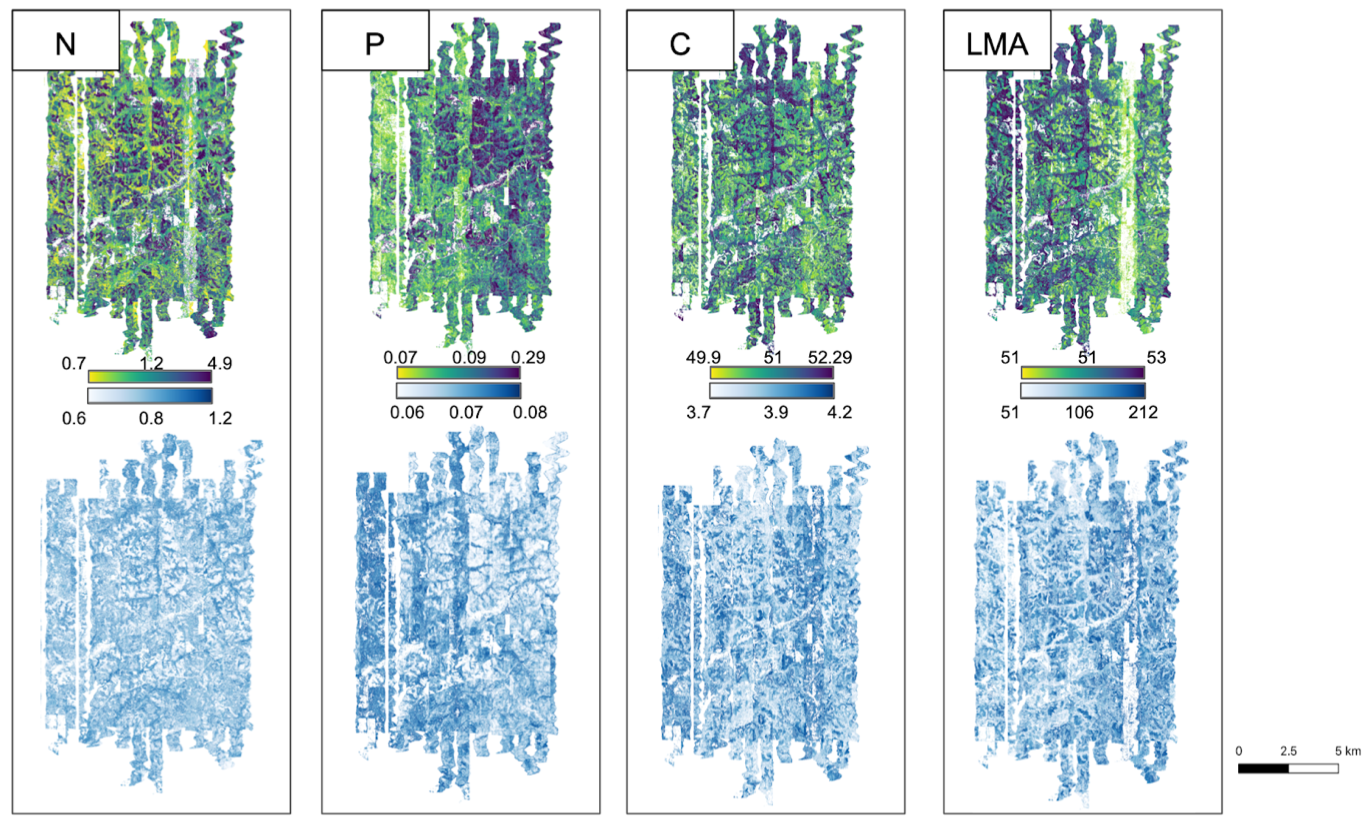

*Figure S. 12. Thematic maps for LMA, %N and %P for the OSBS site (145 km2). The maps show the centroid of around 2.5 million trees, colored by quantile gradient. The top panel (viridis palette) represents individual predictions for %N, %P, %C and LMA. The lower panel (blue) represents the uncertainty for each tree-based prediction, plotted as the width of the 95%PI.*

**Figure S13**

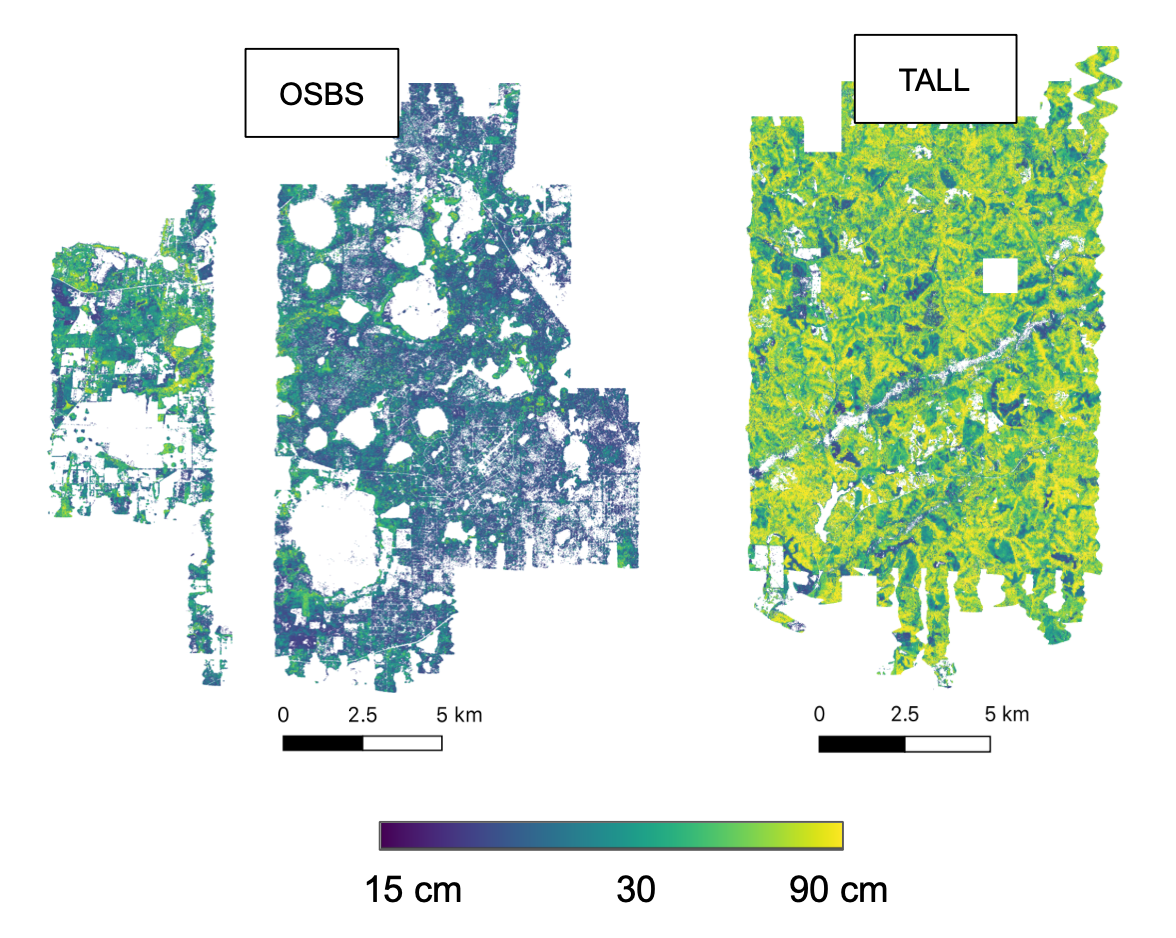

*Figure S. 13 Distribution of BDH (cm) of derived ITCs at TALL and OSBS sites.*

**Figure S14**

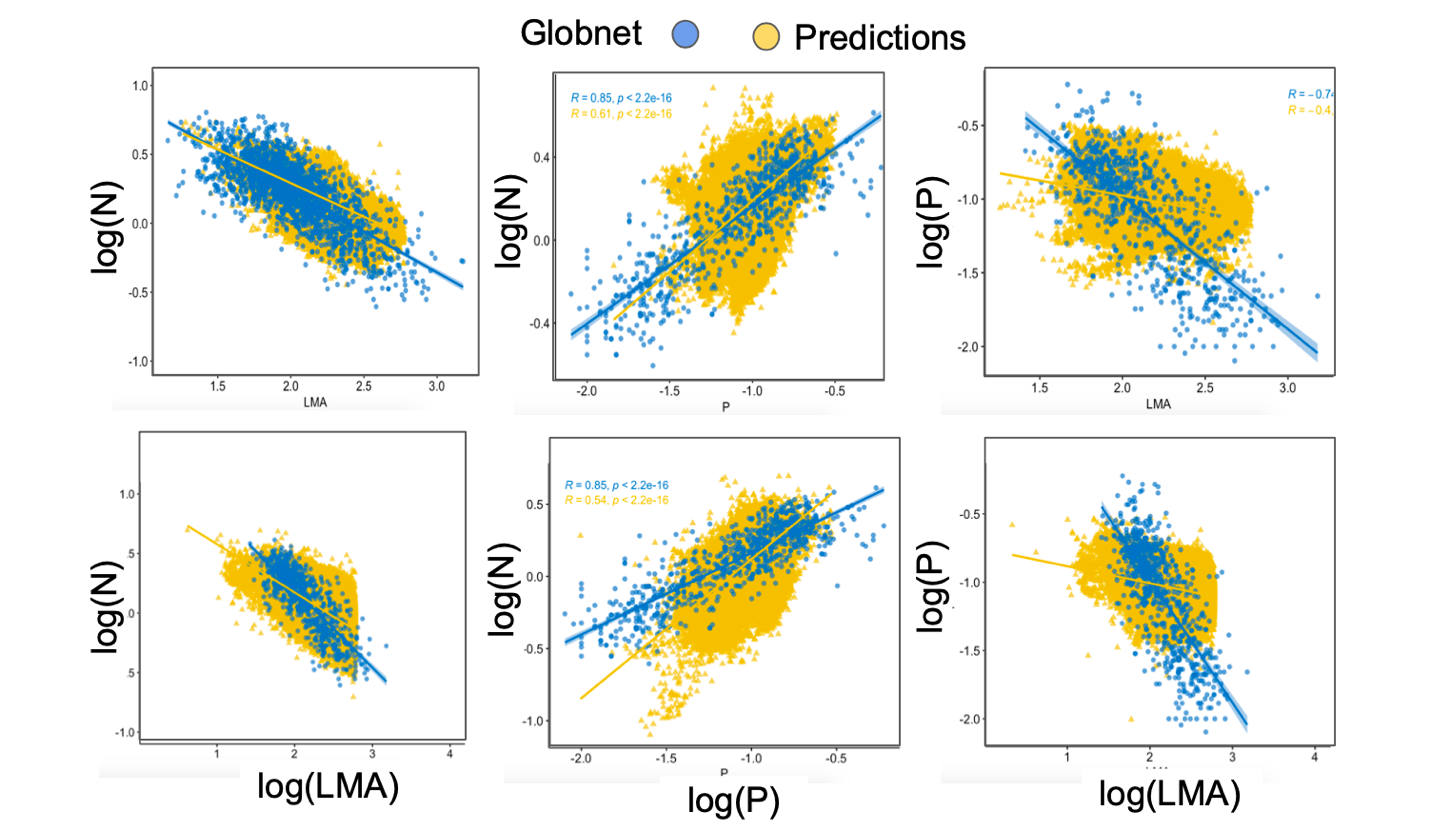

*Figure S.14 Comparison of the leaf economic spectrum relationships between data derived from the pipeline in yellow and data from worldwide leaf economic spectrum dataset in blueTop row shows the log-log relationships in OSBS (N:LMA, N:P, and LMA:P); bottom row shows the log-log relationships in TALL (N:LMA, N:P, and LMA:P)*

**Figure S15**

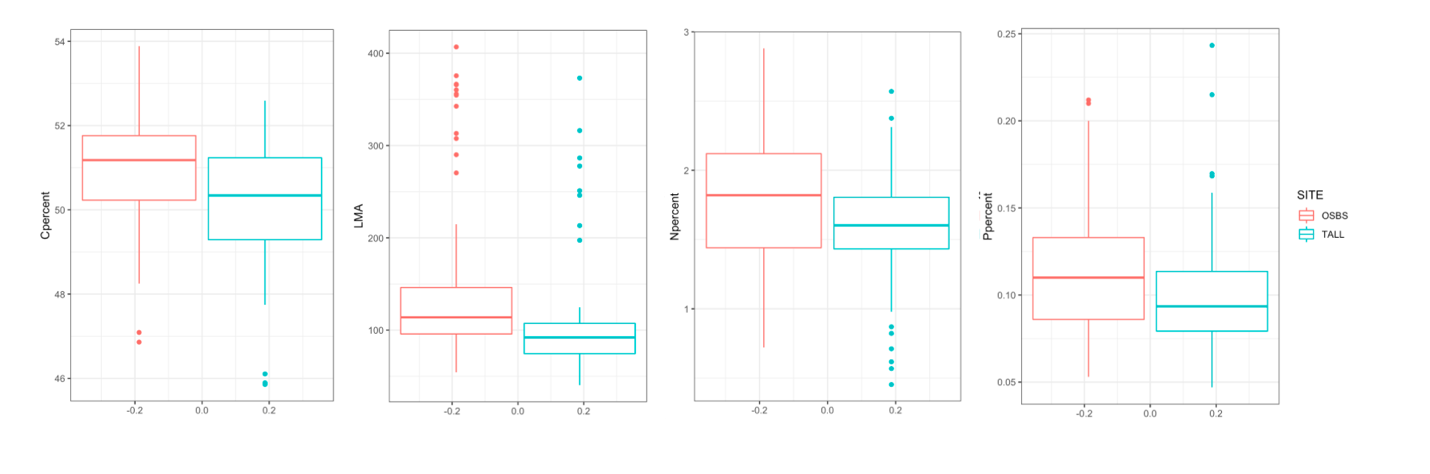

*Figure S.15 Distribution in (a) %C, (B) LMA, (C) %P, and (D) %N across the two sites. On average, OSBS showed higher quantities for all four traits, possibly as an effect of richer soils than in TALL.*

**Figure S16**

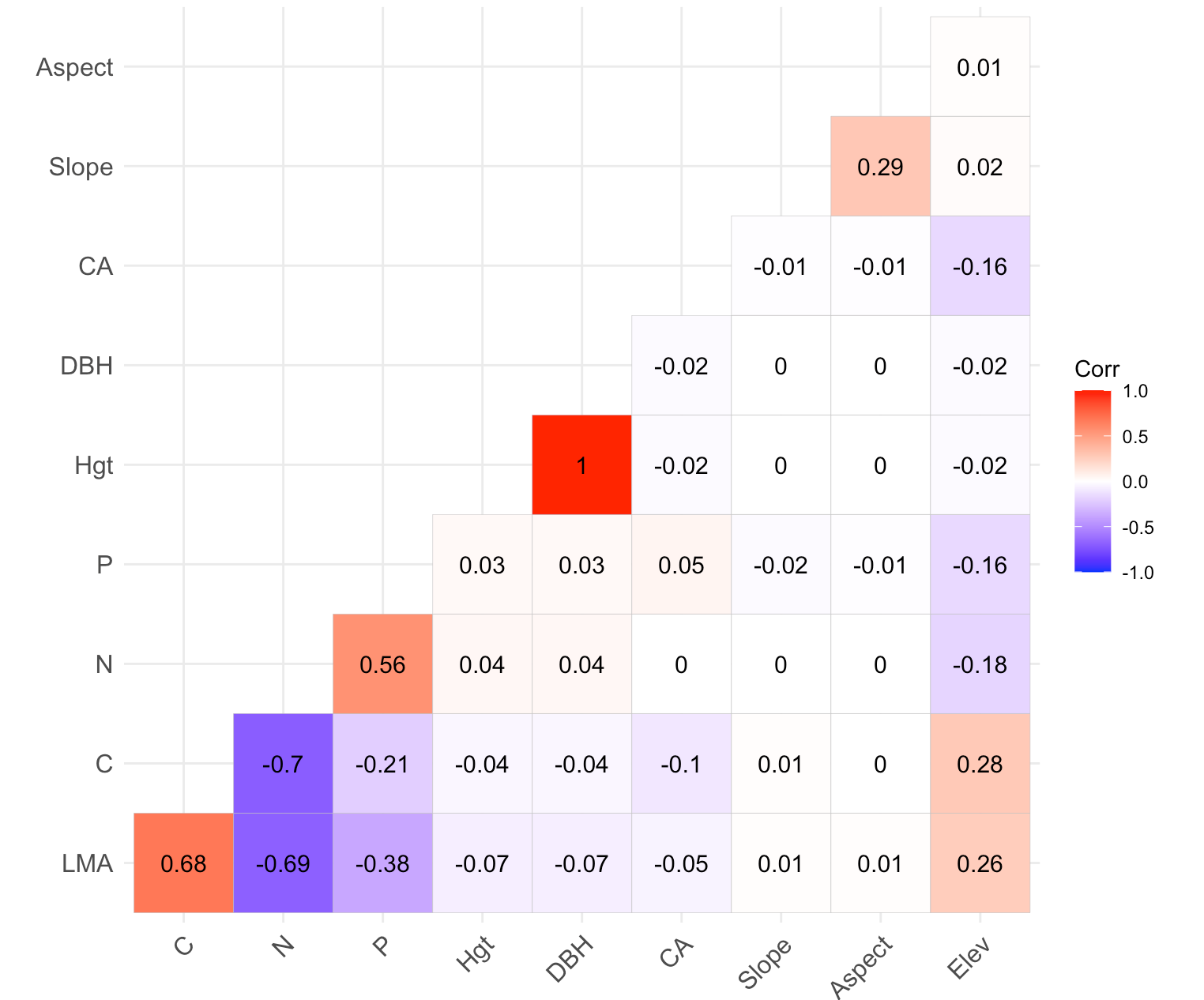

*Figure S. 16. Correlation between terrain, leaf and structural traits for both OSBS and TALL predicted crowns (5M ITCs). Circles size scaled by Pearson’s correlation coefficient. Site specific relationships in appendix.*

**Figure S17**

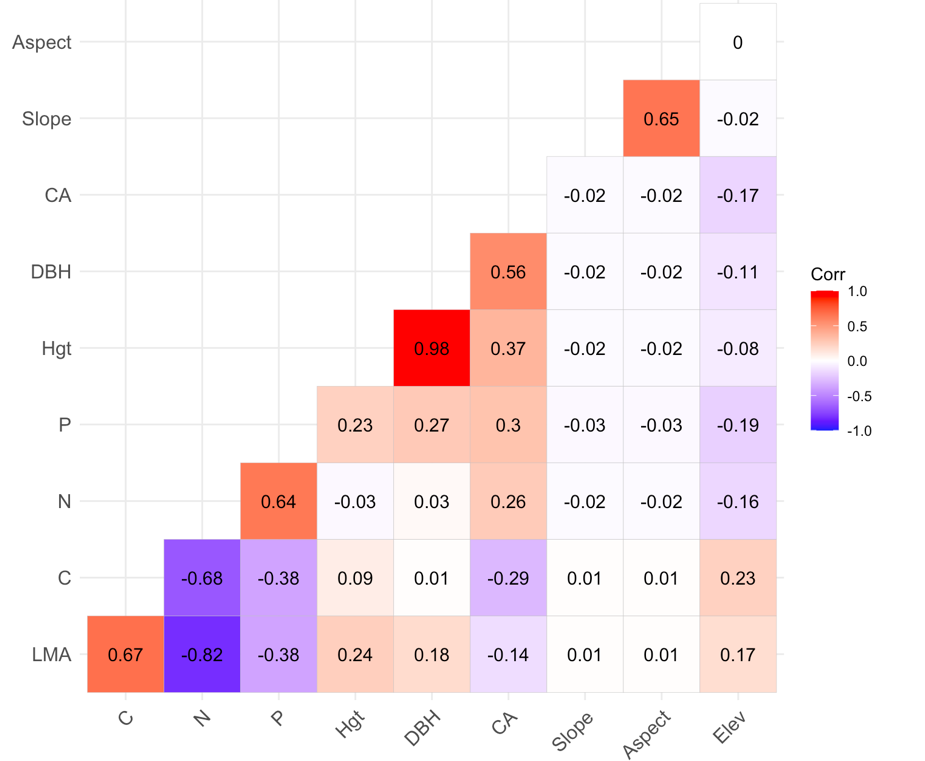

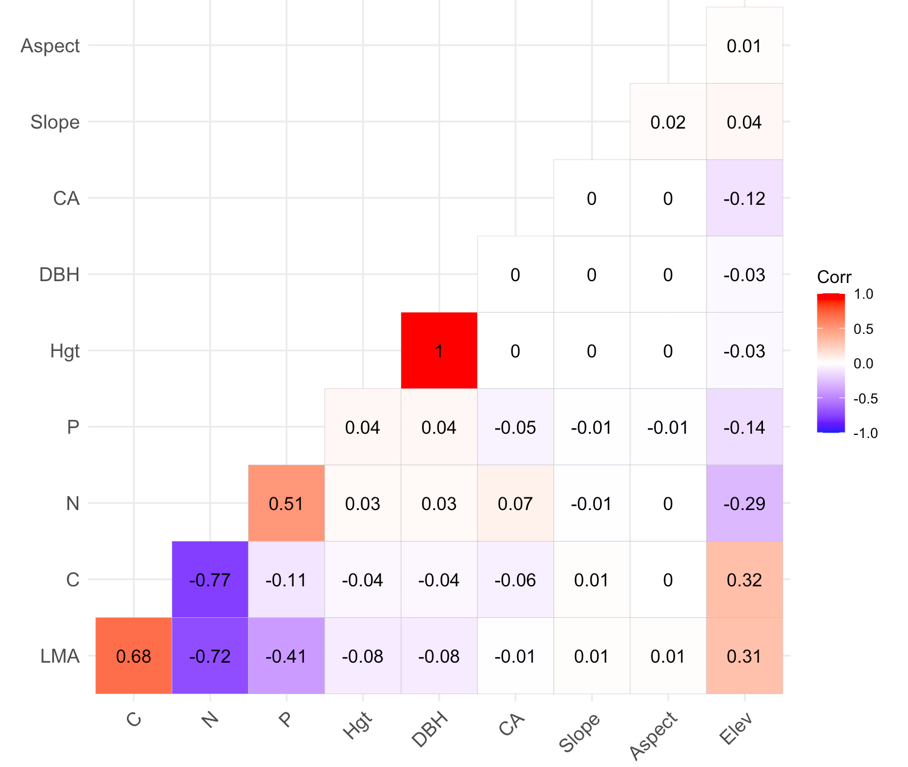

*Figure S.17 Correlation between terrain, leaf and structural traits for OSBS (top) and TALL (bottom) derived tree crowns at landscape scale. Circles size scaled by Pearson’s correlation coefficient.*
